# Albumin alleviates pressure overload-induced heart failure by improving mitochondrial homeostasis via PKA-LKB1-AMPK axis

**DOI:** 10.64898/2026.11.21.768247

**Authors:** Qi An, Boyuan Ma, Yan Fu, Yongzhang Luo

## Abstract

**Aims:** Heart failure (HF) is a global health challenge due to its high incidence, mortality, and socioeconomic burden. Despite medical advancements, HF prognosis remains poor, highlighting the need for enhanced understanding and treatment strategies. Serum albumin, a key circulatory protein, has gained attention as a biomarker in cardiovascular diseases. Low serum albumin levels are linked to HF incidence, progression, and prognosis, suggesting its regulatory role, though mechanisms remain unclear. This study aimed to elucidate the role of serum albumin in HF pathogenesis and investigate its potential therapeutic effects.

**Methods and results:** A mouse model of pressure overload-induced HF was established via transverse aortic constriction (TAC). Serum albumin levels were significantly decreased in HF mice. Administration of recombinant mouse serum albumin (rMSA) effectively restored serum albumin concentrations, improved cardiac systolic function, alleviated cardiac hypertrophy and fibrosis, and reduced inflammatory responses, collectively attenuating disease progression. In contrast, albumin-knockout (*Alb⁻/⁻,* AKO) HF mice displayed aggravated pathological manifestations compared with wild-type (WT) controls, whereas rMSA supplementation markedly ameliorated these impairments. *In vitro* experiments demonstrated that rMSA protected against phenylephrine (PE)-induced hypertrophy in cardiomyocytes. To elucidate the underlying mechanism, transcriptomic analyses of cardiac tissues revealed that albumin played a critical role in improving mitochondrial homeostasis. Experimental validation further demonstrated that albumin activated the PKA-LKB1-AMPK signaling axis, leading to improved mitochondrial respiratory function and oxidative phosphorylation, as well as enhanced mitochondrial quality control. These changes promoted fatty acid oxidation and optimized energy metabolism, thereby reversing the energy imbalance underlying cardiac dysfunction.

**Conclusion:** Our study suggests that serum albumin exerts a protective effect in HF by improving mitochondrial dysfunction and energy metabolism disorders, thus reversing energy imbalance and mitigating disease progression. These results support recombinant serum albumin as a promising therapeutic for HF, with its mechanism involving the critical improvement of mitochondrial homeostasis.

## 1. Introduction

Heart failure (HF), a complex and life-threatening syndrome, has emerged as a significant global public health challenge with escalating prevalence due to aging populations, improved therapies, and enhanced survival rates.^1,2^ The pathogenesis of HF remains incompletely understood, and curative therapies are still limited. Ventricular pressure overload, a common etiology of HF with reduced ejection fraction (HFrEF) and frequently associated with hypertension and aortic stenosis, induces pathological hypertrophy through elevated wall stress, which initially serves as a compensatory mechanism but ultimately progresses to decompensated remodeling involving cardiac systolic dysfunction, hypertrophy, fibrosis, and metabolic dysregulation.^3,4^

HF is fundamentally characterized by an energy supply-demand imbalance, driven by mitochondrial dysfunction and metabolic disorders that disrupt the cardiac high-energy metabolism.^5,6^ Mitochondrial homeostasis, comprising intricate metabolic pathways and dynamic mitochondrial networks, exists in a delicate equilibrium, whereby perturbations induce maladaptive cascades.^7^ Notably, mitochondrial dysfunction is closely linked to HF pathogenesis, with reduced expression and impaired activity of the mitochondrial electron transport chain (ETC) that not only diminish ATP production but also elevate reactive oxygen species (ROS), promoting oxidative damage, cardiomyocyte apoptosis, and pathological ventricular remodeling.^6-9^ Furthermore, ETC dysfunction disrupts mitochondrial quality control (MQC), including biogenesis, fission-fusion dynamics, and mitophagy, further exacerbating cardiac dysfunction.^10,11^ Mitochondrial dysfunction also impairs fatty acid oxidation (FAO) and triggers a pathological metabolic shift in HF.^12,13^ Therefore, restoring mitochondrial homeostasis by targeting mitochondrial dysfunction, metabolic imbalance, and downstream cascades represents a promising therapeutic strategy to reestablish bioenergetic equilibrium in HF.^9,14^

Human serum albumin (HSA), synthesized primarily in the liver, is the most abundant plasma protein.^15,16^ HSA maintains colloid osmotic pressure, binds and transports various ligands, and exhibits antioxidant, anti-inflammatory, anticoagulant, and endothelial regulatory properties.^15-18^ Our recent work demonstrated that administration of young, undamaged recombinant mouse serum albumin (rMSA) not only extends both lifespan and healthspan but also significantly ameliorates type 2 diabetes mellitus in mice.^19,20^ These findings prompt us to investigate whether albumin exerts similar protective effects against HF and to elucidate the underlying mechanisms. Epidemiological studies have long established that hypoalbuminemia (low serum albumin levels, typically < 35 g/L) is an independent predictor of cardiovascular disease incidence, progression, and prognosis, including heart failure irrespective of ejection fraction.^21-30^ Although observational studies have consistently demonstrated an association between low serum albumin levels and adverse HF outcomes, the regulatory roles and mechanisms of albumin in HF progression remain not yet fully elucidated. Crucially, it is still unclear whether therapeutic administration of recombinant serum albumin improves clinical outcomes in HF patients.

This study reveals that reduced serum albumin levels exacerbate pressure overload-induced HF progression. We further establish that albumin can activate the PKA-LKB1-AMPK signaling pathway, which in turn enhances mitochondrial function and fatty acid metabolism. This mechanism effectively improves mitochondrial homeostasis and ameliorates cardiac dysfunction, underscoring the therapeutic potential of recombinant serum albumin for HF.

## 2. Results

### 2.1 Albumin knockout (AKO) aggravates pressure overload-induced heart failure

In this study, pressure overload-induced HF was established in mice by transverse aortic constriction (TAC). Consistent with clinical hypoalbuminemia in HF patients, serum albumin levels were significantly lower in wild type (WT) mice 6 weeks after TAC compared with sham controls (Figure 1A). Hepatic albumin mRNA expression remained unchanged (Figure S1), suggesting altered degradation rather than impaired synthesis in HF.

**Figure 1.**
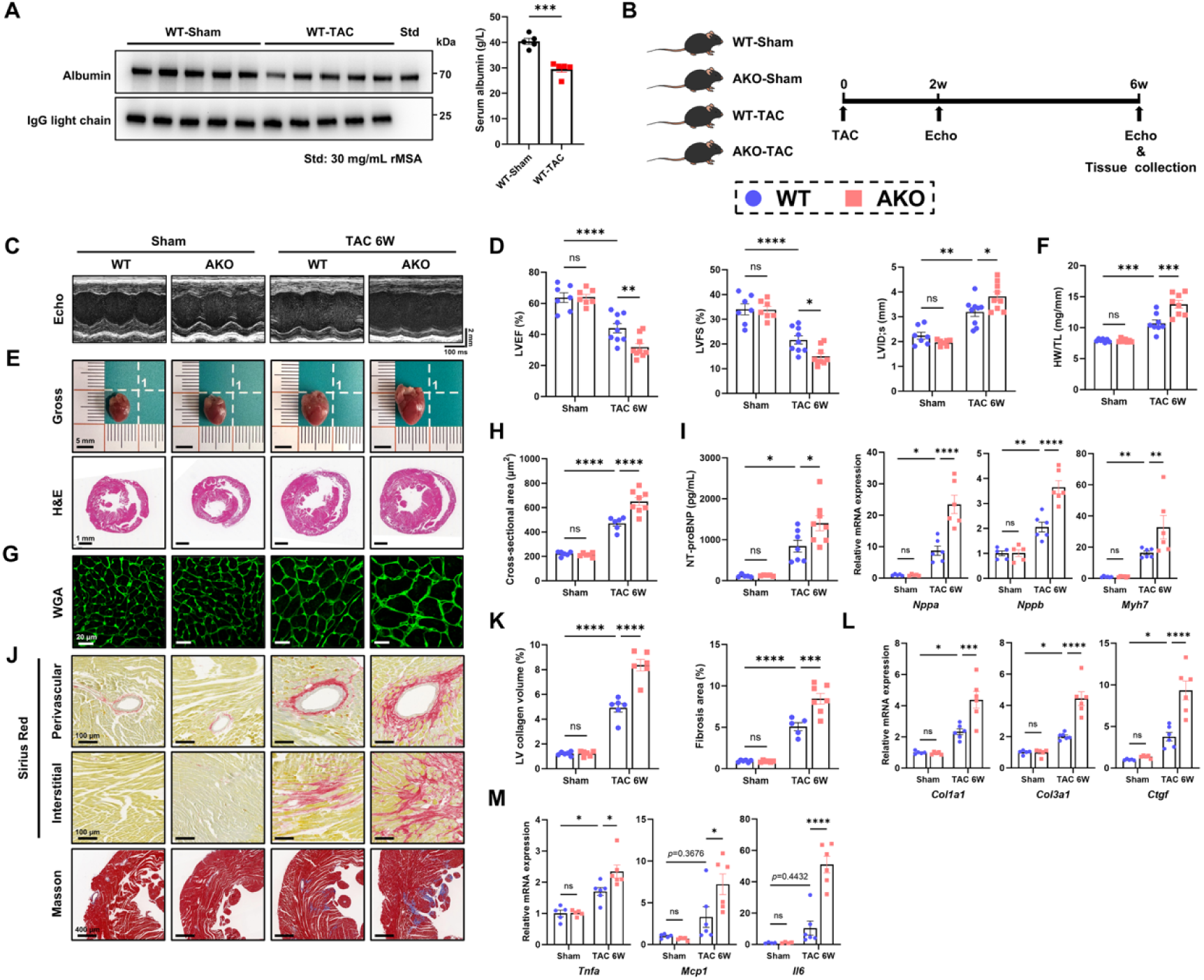
Albumin knockout (AKO) aggravates pressure overload-induced heart failure. (A) Representative immunoblots and quantification of serum albumin levels in WT mice after sham or TAC 6 weeks (n = 5). rMSA standard (Std) (30 mg/mL) is shown as a concentration reference. (B) Experimental design for WT and AKO mice. Echo, echocardiography. (C–M) Analyses were performed in WT and AKO mice after sham or TAC 6 weeks. (C and D) Representative M-mode echocardiographs (C) and quantification (D) of LVEF, LVFS, and LVID;s (n = 7–9). Scale bars, vertical 2 mm and horizontal 100 ms. (E) Representative images of gross morphology (scale bar, 5 mm) and H&E staining (scale bar, 1 mm) of heart sections. (F) Quantification of HW/TL (n = 7–8). (G) Representative images of WGA staining (scale bar, 20 μm) of heart sections. (H) Quantification of cross-sectional area of cardiomyocytes (n = 6–8). (I) Assessment of heart failure markers: serum NT-proBNP levels (n = 5–8) and cardiac mRNA expression of *Nppa*, *Nppb*, and *Myh7* (n = 5–6). (J) Representative images of perivascular and interstitial Sirius red staining (scale bar, 100 μm) and Masson’s trichrome staining (scale bar, 400 μm) of heart sections. (K) Quantification of LV collagen volume fraction (Sirius red) (n = 6) and fibrosis area fraction (Masson’s trichrome) (n = 5–7) of heart sections. (L) Cardiac mRNA expression of fibrotic genes (n = 5–6). (M) Cardiac mRNA expression of inflammatory genes (n = 5–6). (I, L and M) mRNA data normalized to *Actb* mRNA. Mean ± SEM, two-way ANOVA followed by Tukey’s multiple comparisons test (D, F, H, I, K–M) or using unpaired two-tailed Student’s *t*-test (A). ns, not significant; \**p*<0.05, \*\**p*<0.01, \*\*\**p*<0.001, \*\*\*\**p*<0.0001.

To explore the specific role of albumin in cardiac function, we used albumin knockout (*Alb*^-/-^, AKO) mice, which exhibit severe albumin deficiency as an extreme pathological state. Serum analysis confirmed the complete absence of albumin expression in AKO mice versus WT (Figure S2A). To investigate the intrinsic cardiac effects of albumin deficiency, we first compared cardiac function between AKO and WT mice at baseline across different age groups. Echocardiography revealed no significant differences in left ventricular ejection fraction (LVEF), fractional shortening (LVFS), end-systolic internal dimensions (LVID;s), or corrected LV mass (LV mass cor) between AKO and WT mice at either 2 or 24 months of age, while both WT and AKO mice exhibited evident age-related impairment in left ventricular systolic function when 24-month-old animals were compared with their 2-month-old counterparts (Figure S2B and C, Table S1). Heart weight/tibia length ratios (HW/TL) and serum NT-proBNP levels increased with age but did not differ by genotype (Figure S2D and E). Thus, genetic albumin deficiency alone did not exacerbate cardiac dysfunction or remodeling in both young and aged mice without any external intervention.

To investigate the role of albumin under pathological stress, we subjected WT and AKO mice to pressure overload via TAC to induce HF (Figure 1B). At 2 weeks post-TAC, AKO-TAC mice showed more severe LVEF decline and LVID;s elevation than WT-TAC mice on echocardiography (Figure S3 and Table S2). By 6 weeks, this dysfunction advanced, with AKO-TAC mice exhibiting additional deterioration in LVEF, LVFS, and LVID;s compared to WT-TAC controls (Figure 1C and D, Table S3). AKO-TAC mice also displayed increased HW/TL ratios, with gross morphology and H&E staining revealing exacerbated cardiac hypertrophy (Figure 1E and F). Furthermore, cardiomyocyte cross-sectional areas were significantly larger in AKO-TAC than in WT-TAC mice (Figure 1G and H). At 6 weeks post-TAC, serum NT-proBNP levels and cardiac mRNA levels of remodeling markers (*Nppa*, *Nppb*, *Myh7*) were significantly elevated in AKO-TAC mice (Figure 1I). Moreover, fibrosis was more severe in AKO-TAC hearts, as shown by increased interstitial and perivascular collagen deposition in Masson’s trichrome and Sirius red staining (Figure 1J and K), along with upregulation of fibrotic genes (*Col1a1*, *Col3a1*, *Ctgf*) (Figure 1L). Additionally, cardiac mRNA expression of proinflammatory cytokines (*Tnfa*, *Mcp1*, *Il6*) was significantly higher in AKO-TAC mice (Figure 1M). Notably, no differences were observed between WT-Sham and AKO-Sham groups in any of these measures (Figure 1C–M).

Together, these findings demonstrate that albumin deficiency exacerbates cardiac dysfunction, hypertrophy, fibrosis, and inflammation, worsening pathological remodeling in pressure overload-induced HF.

### 2.2 Administration of rMSA alleviates pressure overload-induced heart failure in WT mice or mitigates disease progression in AKO mice

Given the aggravated HF phenotype in AKO mice, we hypothesized that albumin could play a protective role and that its supplementation could mitigate disease progression by elevating serum albumin levels. To test this, recombinant mouse serum albumin (rMSA) or saline was administered intravenously to WT mice before and after TAC surgery (Figure 2A). Prior studies established the safety and efficacy of this rMSA dosage in mice, with no observed toxicity during chronic administration.^19,20^ At 8 weeks post-TAC, serum albumin levels were restored in WT-rMSA-TAC mice compared with WT-Saline-TAC controls with lower serum albumin levels (Figure 2B). rMSA treatment improved systolic function, as shown by higher LVEF and LVFS alongside reduced LVIDs (Figure 2C and D, Table S4). Cardiac hypertrophy was attenuated, with lower HW/TL ratios and smaller cardiomyocyte cross-sectional areas in WT-rMSA-TAC mice (Figure 2E–H). Serum NT-proBNP and cardiac mRNA levels of remodeling markers (*Nppa*, *Nppb*, *Myh7*) were also reduced (Figure 2I). Furthermore, interstitial and perivascular fibrosis were less severe in WT-rMSA-TAC hearts, accompanied by downregulated expression of fibrotic genes (*Col1a1*, *Col3a1*, *Ctgf*, *Tgfb1*) (Figure 2J– L). In addition, the cardiac inflammatory response was suppressed, with lower mRNA levels of *Tnfa* and *Il6* (Figure 2M). No significant differences were observed between sham groups (Figure 2C–M).

**Figure 2.**
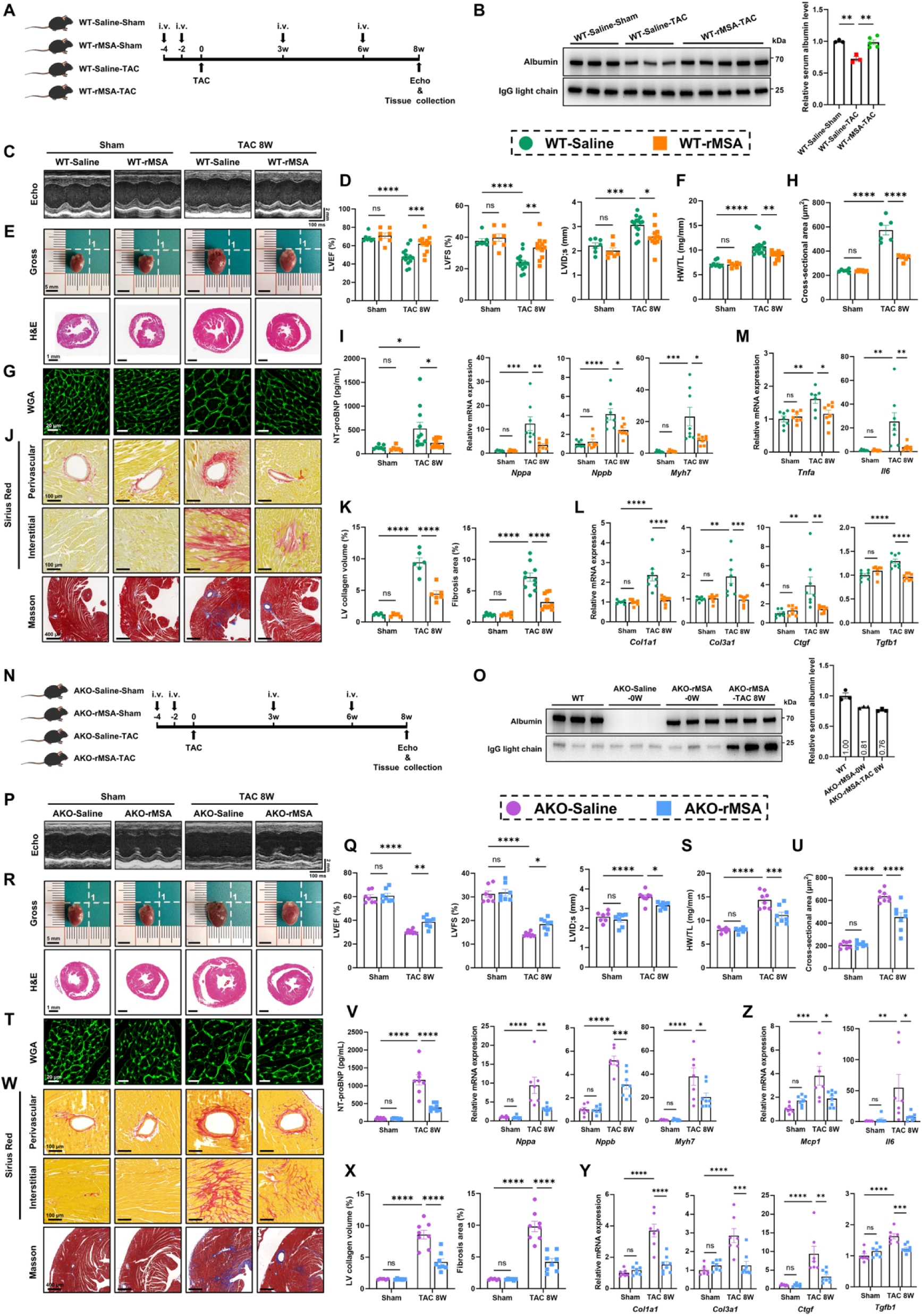
Administration of rMSA alleviates pressure overload-induced heart failure in WT mice or mitigates disease progression in AKO mice. (A) Experimental design for WT mice. Mice received intravenous administration of rMSA at a dosage of 1.5 mg/g or an equivalent volume of sterile saline. Echo, echocardiography; i.v., intravenous injection. (B) Representative immunoblots and quantification of relative serum albumin levels in WT-Saline-Sham, WT-Saline-TAC and WT-rMSA-TAC mice after sham or TAC 8 weeks (n = 3–5). (C–M) Analyses were performed in WT mice treated with rMSA or saline after sham or TAC 8 weeks. (C and D) Representative M-mode echocardiographs (C) and quantification (D) of LVEF, LVFS, and LVID;s in WT mice (n = 7–14). Scale bars, vertical 2 mm and horizontal 100 ms. (E) Representative images of gross morphology (scale bar, 5 mm) and H&E staining (scale bar, 1 mm) of heart sections in WT mice (F) Quantification of HW/TL in WT mice (n = 7–14). (G) Representative images of WGA staining (scale bar, 20 μm) of heart sections in WT mice. (H) Quantification of cross-sectional area of cardiomyocytes in WT mice (n = 6). (I) Assessment of heart failure markers: serum NT-proBNP levels (n = 6–11), and cardiac mRNA expression of *Nppa*, *Nppb* and *Myh7* (n = 7–8) in WT mice. (J) Representative images of perivascular and interstitial Sirius red staining (scale bar, 100 μm) and Masson’s trichrome staining (scale bar, 400 μm) of heart sections in WT mice. (K) Quantification of LV collagen volume fraction (Sirius red) of heart sections (n = 5– 6), and fibrosis area fraction (Masson’s trichrome) of heart sections (n = 6–10) in WT mice. (L) Cardiac mRNA expression of fibrotic genes in WT mice (n = 7–8). (M) Cardiac mRNA expression of inflammatory genes in WT mice (n = 7–8). (N) Experimental design for AKO mice. Mice received intravenous administration of rMSA at a dosage of 1.5 mg/g or an equivalent volume of sterile saline. Echo, echocardiography; i.v., intravenous injection. (O) Representative immunoblots and quantification of relative serum albumin levels in saline-or rMSA-treated AKO mice, assessed pre-TAC (0W) and post-TAC (8W), compared to WT controls (n = 3). (P–Z) Analyses were performed in AKO mice treated with rMSA or saline after sham or TAC 8 weeks. (P and Q) Representative M-mode echocardiographs (P) and quantification (Q) of LVEF, LVFS, and LVID;s in AKO mice (n = 7–8). Scale bars, vertical 2 mm and horizontal 100 ms. (R) Representative images of gross morphology (scale bar, 5 mm) and H&E staining (scale bar, 1 mm) of heart sections in AKO mice. (S) Quantification of HW/TL in AKO mice (n = 7–8). (T) Representative images of WGA staining (scale bar, 20 μm) of heart sections in AKO mice. (U) Quantification of cross-sectional area of cardiomyocytes in AKO mice (n = 7–8). (V) Assessment of heart failure markers: serum NT-proBNP levels (n = 7–8), and cardiac mRNA expression of *Nppa*, *Nppb* and *Myh7* (n = 7–8) in AKO mice. (W) Representative images of perivascular and interstitial Sirius red staining (scale bar, 100 μm) and Masson’s trichrome staining (scale bar, 400 μm) of heart sections in AKO mice. (X) Quantification of LV collagen volume fraction (Sirius red) of heart sections (n = 6– 8), and fibrosis area fraction (Masson’s trichrome) of heart sections (n = 6–8) in AKO mice. (Y) Cardiac mRNA expression of fibrotic genes in AKO mice (n = 7–8). (Z) Cardiac mRNA expression of inflammatory genes in AKO mice (n = 7–8). (I, L, M, V, Y and Z) mRNA data normalized to *Actb* mRNA. Mean ± SEM, two-way ANOVA followed by Tukey’s multiple comparisons test. ns, not significant; \**p*<0.05, \*\**p*<0.01, \*\*\**p*<0.001, \*\*\*\**p*<0.0001.

To determine whether the protective effect is mediated specifically by circulating albumin, we treated AKO mice with rMSA to restore circulating albumin levels and evaluate therapeutic efficacy. The administration regimen was consistent with that described above (Figure 2N). We confirmed that rMSA administration in AKO mice restored serum albumin levels to approximately 80% of those in WT controls, both before surgery and at the experimental endpoint (Figure 2O). rMSA administration significantly improved cardiac function in AKO-rMSA-TAC mice, evidenced by increased LVEF and LVFS, alongside reduced LVIDs (Figure 2P and Q, Table S5). Cardiac hypertrophy was also attenuated, as shown by decreased HW/TL ratios and smaller cardiomyocyte cross-sectional areas (Figure 2R–U). At the molecular level, serum NT-proBNP levels and cardiac expression of remodeling markers (*Nppa*, *Nppb*, *Myh7*) were downregulated in AKO-rMSA-TAC mice (Figure 2V). Furthermore, rMSA treatment alleviated fibrosis, with reduced collagen deposition and downregulated expression of fibrotic genes (Figure 2W–Y), and suppressed cardiac inflammation, indicated by lower mRNA levels of *Mcp1* and *Il6* (Figure 2Z). No significant differences were observed between sham groups in any of these parameters (Figure 2P– Z). These data reveal that exogenous rMSA mitigates disease progression in AKO mice with pressure overload-induced HF.

Collectively, it is evident that exogenous rMSA supplementation restores serum albumin levels and ameliorates pressure overload-induced HF, thereby highlighting the therapeutic potential of albumin administration in clinical HF.

### 2.3 RNA sequencing reveals that albumin ameliorates pressure overload-induced cardiac mitochondrial dysfunction and fatty acid metabolism disorders

To elucidate the mechanism by which albumin alleviates HF, we performed transcriptomic profiling on cardiac tissues from WT-Sham, AKO-Sham, WT-TAC, and AKO-TAC mice. Under baseline conditions, AKO-Sham and WT-Sham hearts showed minimal transcriptional differences with 175 differentially expressed genes (DEGs), consistent with their comparable cardiac phenotype (Figure S4A). In contrast, profound transcriptional dysregulation was observed in AKO-TAC versus WT-TAC hearts, with 1525 genes upregulated and 1353 downregulated (Figure S4B). Upregulated genes in AKO-TAC hearts were enriched in pathways related to extracellular matrix remodeling and inflammatory responses based on Gene Ontology (GO) analysis (Figure S4C), aligning with the observed exacerbated fibrosis and inflammation. Conversely, downregulated genes were strongly clustered in mitochondrial energy metabolism pathways, including oxidative phosphorylation (OXPHOS), tricarboxylic acid (TCA) cycle, fatty acid metabolism, and respiratory electron transport, as shown by GO and KEGG analyses (Figure 3A, S4D and E). Heatmap visualization confirmed a severe suppression of genes critical for mitochondrial function and fatty acid metabolism in AKO-TAC hearts compared to WT-TAC controls (Figure S4F and G). This suppression was also present, though less severe, in WT-TAC versus WT-Sham hearts, reflecting the hallmark energy metabolic impairment in HF (Figure S4F and G). GSEA further highlighted the divergent biological processes: AKO-TAC hearts were enriched for pro-fibrotic and pro-inflammatory processes, whereas WT-TAC hearts retained greater enrichment for mitochondrial and metabolic pathways, as shown in the key enrichment plots (Figure S4H and I).

**Figure 3.**
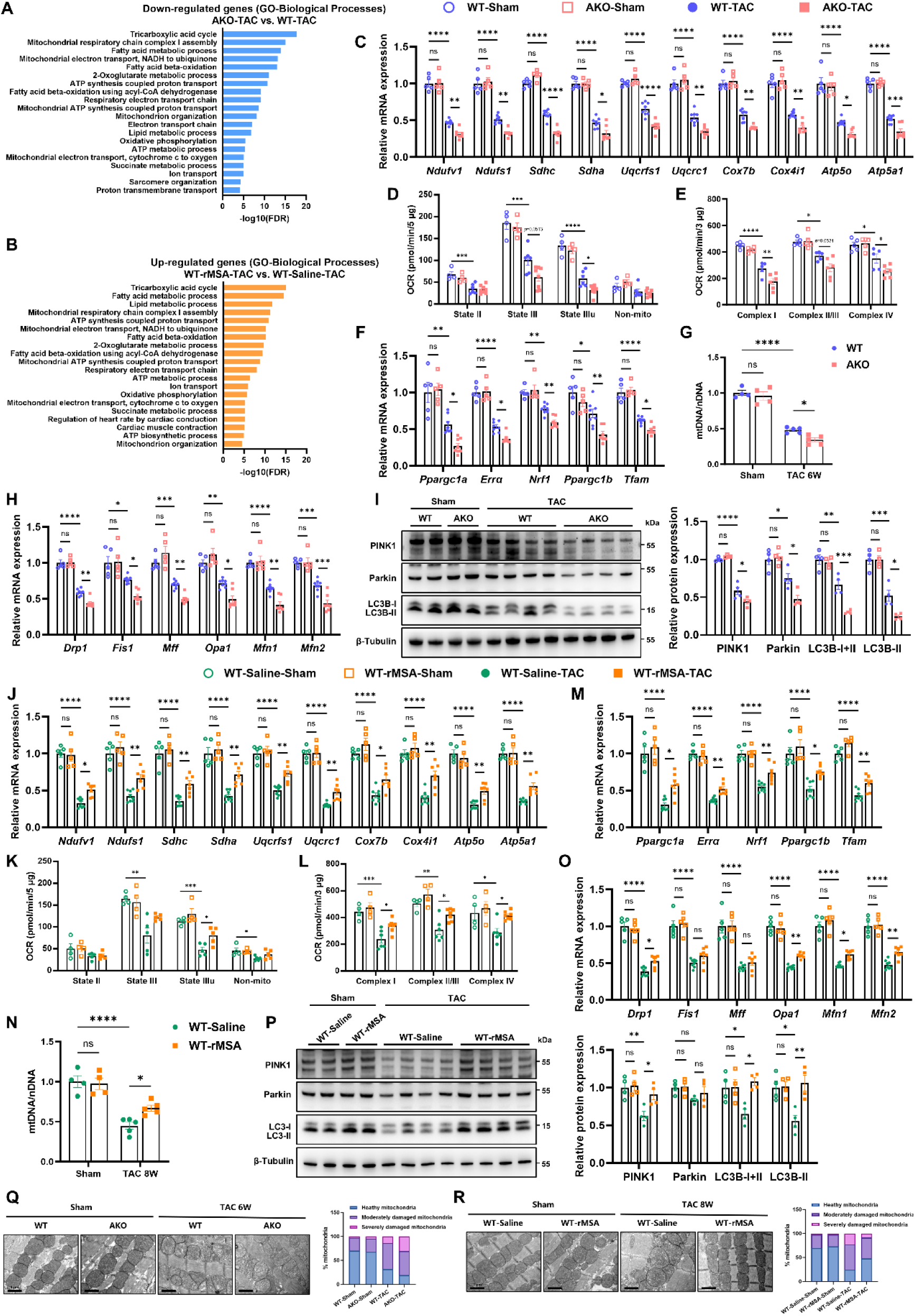
Albumin improves cardiac mitochondrial respiratory function and maintains cardiac mitochondrial quality control in pressure overload-induced heart failure. (A–B) GO enrichment analysis of biological processes for genes down-regulated in AKO-TAC hearts (vs. WT-TAC) (A, n=4) or up-regulated in WT-rMSA-TAC hearts (vs. WT-Saline-TAC) (B, n=3). Representative enriched terms (FDR < 0.05) were ranked by significance (-log₁₀FDR). FDR, false discovery rate (*Q* value). (C–I) Samples from WT and AKO mice after sham or TAC 6 weeks. (C) Cardiac mRNA expression of ETC genes (n = 5–8). (D and E) Quantification of the coupling assay (D, n = 4–8), and the electron flow assay (E, n = 5–6) in isolated cardiac mitochondria. See measurement plots in Figure S8. (F) Cardiac mRNA expression of mitochondrial biogenesis-associated genes (n = 5–8). (G) mtDNA/nDNA ratio in cardiac tissues (n = 4–5). (H) Cardiac mRNA expression of mitochondrial fission-and fusion-associated genes (n = 5–8). (I) Representative immunoblots and quantitation of mitophagy-associated proteins in cardiac tissues (n = 4). (J–P) Samples from WT mice treated with rMSA or saline after sham or TAC 8 weeks. (J) Cardiac mRNA expression of ETC genes (n = 5–8). (K and L) Quantification of the coupling assay (K, n = 4–5), and the electron flow assay of isolated cardiac mitochondria (L, n = 4–7). See measurement plots in Figure S8. (M) Cardiac mRNA expression of mitochondrial biogenesis-associated genes (n = 5– 8). (N) mtDNA/nDNA ratio in cardiac tissues (n = 4–5). (O) mRNA expression of mitochondrial fission- and fusion-associated genes (n = 5–8). (P) Representative immunoblots and quantitation of mitophagy-associated proteins in cardiac tissues (n = 4). (Q and R) Representative TEM images (scale bar, 1 μm) and quantitative classification of mitochondrial morphology in cardiac tissues from WT and AKO mice after sham or TAC 6 weeks (Q, n = 4), and from WT mice treated with rMSA or saline after sham or TAC 8 weeks (R, n = 4). Mitochondria from random fields (>200 mitochondria per mouse) were classified as healthy, moderately damaged, or severely damaged. ETC genes including *Ndufv1* and *Ndufs1* (Complex I), *Sdhc* and *Sdha* (Complex II), *Uqcrfs1* and *Uqcrc1* (Complex III), *Cox7b* and *Cox4i1* (Complex IV), and *Atp5o* and *Atp5a1* (Complex V) were detected in (C) and (J); Mitochondrial biogenesis-associated genes including *Ppargc1a*, *Errα*, *Nrf1*, *Ppargc1b* and *Tfam* were detected in (F) and (M); Mitochondrial fission-associated genes including *Drp1*, *Fis1* and *Mff*, and fusion-associated genes including *Opa1*, *Mfn1* and *Mfn2* were detected in (H) and (O). mRNA data normalized to *Actb* mRNA. Mitophagy-associated proteins including PINK1, Parkin, and LC3B-I/II were detected in (I) and (P), and β-Tubulin served as the loading control. Mean ± SEM, two-way ANOVA followed by Tukey’s multiple comparisons test. ns, not significant; \**p*<0.05, \*\**p*<0.01, \*\*\**p*<0.001, \*\*\*\**p*<0.0001. mtDNA, mitochondrial DNA; nDNA, nuclear DNA.

Transcriptomic analysis of rMSA-treated hearts (WT-rMSA-TAC vs. WT-Saline-TAC) revealed an opposite pattern, with 1297 genes upregulated and 1464 downregulated (Figure S5A). GO analysis demonstrated that the upregulated genes in WT-rMSA-TAC hearts were enriched in mitochondrial metabolic pathways, such as the TCA cycle, fatty acid β-oxidation, OXPHOS, and respiratory electron transport, which was corroborated by KEGG pathway analysis (Figure 3B, S5B and C). Notably, these enriched processes were suppressed in AKO-TAC hearts, suggesting that albumin may help restore cardiac mitochondrial homeostasis. Heatmaps further showed enhanced expression of mitochondrial metabolic genes in WT-rMSA-TAC versus WT-Saline-TAC hearts (Figure S5D and E). Meanwhile, genes downregulated by rMSA treatment were involved in extracellular matrix organization and inflammatory responses (Figure S5F), pathways that were conversely upregulated in AKO-TAC hearts. GSEA further highlighted the preferential enrichment of mitochondrial bioenergetic pathways in WT-rMSA-TAC hearts, as shown in the key enrichment plots (Figure S5G and H).

Integrated transcriptomic analysis reveals that albumin deficiency under pressure overload is associated with severe disruption of genes involved in cardiac mitochondrial energy metabolism and fatty acid utilization. The restoration of these pathways upon exogenous albumin administration supports the inference that albumin may protect against HF, in part, by preserving these vital metabolic processes.

Immunofluorescence staining of cardiac tissue sections confirmed albumin localization in cardiomyocytes, with specific signals markedly stronger than those of secondary antibody-only controls (Figure S6A). In primary neonatal mouse cardiomyocytes (NMCMs) incubated with rMSA, immunofluorescence further validated albumin internalization into cardiomyocytes (Figure S6B). Considering the cellular uptake of albumin and the pivotal role of mitochondrial dysfunction in HF, we proposed that albumin conferred direct cardioprotection via regulating cardiomyocyte energetics, and attenuated HF progression by maintaining cardiomyocyte mitochondrial homeostasis.

### 2.4 Albumin improves cardiac mitochondrial respiratory function and maintains cardiac mitochondrial quality control in pressure overload-induced heart failure

Mitochondrial dysfunction, a hallmark of HF, disrupts ATP production and energy homeostasis. Clinical and animal studies corroborate this association, showing repressed OXPHOS-related gene expression and diminished OXPHOS capacity in HF hearts.^8,31^ Notably, our transcriptomic analysis indicated significant downregulation of mitochondrial electron transport chain (ETC) genes in AKO-TAC hearts compared to WT-TAC controls (Figure S4F, G and I). This was corroborated by qRT-PCR and immunoblotting, which revealed a marked TAC-induced reduction in ETC complex expression that was further exacerbated in AKO-TAC hearts (Figure 3C and S7A). For further evaluation of mitochondrial respiratory capacity, cardiac tissue mitochondria were isolated and subjected to Seahorse functional assays. The coupling assay measuring complex I-driven respiration showed that State III and State IIIu respiration were significantly lower in WT-TAC versus WT-Sham mitochondria and declined further in AKO-TAC mitochondria (Figure 3D and S7B). The electron flow assay assessing ETC efficiency also demonstrated severely impaired activities of ETC complexes I, II/III, and IV in AKO-TAC mitochondria compared to WT-TAC (Figure 3E and S7C).

Beyond enhancing mitochondrial respiratory capacity, albumin may also sustain cardiac mitochondrial quality control (MQC) in pressure overload-induced HF. Our transcriptomic data highlighted “mitochondrion organization” as a key biological process disrupted by albumin deficiency and restored by rMSA administration (Figure S4F and S5D). MQC plays a pivotal role in HF by maintaining structural and functional mitochondrial integrity through selective clearance of impaired organelles, thus preventing cardiomyocyte dysfunction.^6,11^ qRT-PCR and immunoblotting showed significant downregulation of core mitochondrial biogenesis factors (PGC-1α, ERRα, NRF1, PGC-1β, TFAM) in WT-TAC hearts versus controls, with further suppression in AKO-TAC hearts (Figure 3F and S7D). AKO-TAC hearts also exhibited lower mitochondrial genomic content (mtDNA/nDNA ratio) relative to WT-TAC controls (Figure 3G). Moreover, qRT-PCR revealed TAC-induced downregulation of key mitochondrial dynamics genes in WT-TAC versus WT-Sham hearts, including fission-associated genes (*Drp1*, *Fis1*, and *Mff*) and fusion-associated genes (*Opa1*, *Mfn1*, and *Mfn2*), with worsened suppression in AKO-TAC hearts (Figure 3H). Concurrently, immunoblotting demonstrated reduced mitophagy-related protein levels (PINK1, Parkin, LC3B) in WT-TAC versus WT-Sham controls, with more marked decreases in AKO-TAC hearts, and no baseline differences existed between sham groups (Figure 3I).

Exogenous albumin administration reversed these deficits. Transcriptomic analysis revealed significant enrichment of OXPHOS and mitochondrial respiratory chain pathways in WT-rMSA-TAC hearts compared to WT-Saline-TAC controls (Figure S5D, E and H). Accordingly, we confirmed that rMSA treatment attenuated the TAC-induced downregulation of ETC genes and proteins in WT mice (Figure 3J and S7E). Functionally, the coupling assay showed reduced State III and State IIIu OCR in WT-Saline-TAC mitochondria relative to WT-Saline-Sham controls, while rMSA restored State IIIu OCR in WT-rMSA-TAC mitochondria (Figure 3K and S7F). In addition, the electron flow assay further confirmed that rMSA enhanced complex I-, II/III-, and IV-driven respiration in WT-rMSA-TAC mitochondria (Figure 3L and S7G). Furthermore, rMSA treatment enhanced the MQC mechanism. In TAC-induced WT mice, rMSA upregulated mitochondrial biogenesis genes and proteins (Figure 3M and S7H), increased mtDNA/nDNA ratio (Figure 3N), enhanced fusion gene expression (Figure 3O), and elevated mitophagy protein levels in hearts relative to WT-Saline-Sham controls, and no baseline differences existed between sham groups (Figure 3P).

Mitochondrial ultrastructural abnormalities, including cristae disorganization, fragmentation, swelling, and abnormal matrix remodeling, are hallmark features of HF progression, and drive a vicious cycle that exacerbates cardiac metabolic dysfunction and disease progression.^6^ To assess albumin’s effects on mitochondrial ultrastructure, we analyzed cardiac mitochondria from experimental mice via transmission electron microscopy (TEM), categorizing them into three morphological types: healthy, moderately damaged, and severely damaged. AKO-TAC hearts contained more severely damaged mitochondria than WT-TAC hearts, and rMSA treatment reduced the proportion of abnormal mitochondria in TAC-induced WT mice, as WT-rMSA-TAC mice exhibited less mitochondrial swelling and cristae disruption versus WT-Saline-TAC controls (Figure 3Q and R).

To evaluate the rescue efficacy of rMSA in TAC-induced AKO mice, parallel assays were performed. rMSA administration partially rescued the expression of ETC components (Figure S8A and B), improved State III/IIIu respiration in the coupling assay (Figure S8C), and restored complex I-, II/III-, and IV-driven respiration in the electron flow assay in TAC-induced AKO mice (Figure S8D). Additionally, rMSA administration restored the MQC mechanism in TAC-induced AKO mice, as evidenced by upregulated expression of mitochondrial biogenesis genes and proteins (Figure S8E and F), along with increased mtDNA/nDNA ratio in AKO-rMSA-TAC hearts compared to AKO-Saline-TAC controls (Figure S8G). Furthermore, rMSA treatment elevated mitochondrial fusion and fission gene expression (Figure S8H), and mitophagy protein levels in hearts relative to AKO-Saline-Sham controls (Figure S8I). TEM analysis revealed that cardiomyocyte mitochondrial integrity was better preserved in AKO-rMSA-TAC mice than in AKO-Saline-TAC controls, as indicated by less swelling and cristae disruption (Figure S8J). No differences were observed between corresponding sham groups.

Moreover, in isolated primary NMCMs with or without rMSA treatment, qRT-PCR revealed that most MQC-related genes were significantly upregulated in rMSA-treated cardiomyocytes relative to controls (Figure S9).

Taken together, these results indicate that albumin deficiency exacerbates pressure overload-induced mitochondrial respiratory dysfunction, impairs MQC, and disrupts mitochondrial ultrastructural integrity. Conversely, albumin supplementation can preserve these mitochondrial properties, thereby supporting cardiac metabolic homeostasis and energy supply in HF.

### 2.5 Albumin enhances cardiac fatty acid metabolism

Cardiac metabolism in HF shifts from fatty acid oxidation (FAO) dominance toward impaired pathways.^12,32^ Transcriptomics revealed severe suppression of FAO-related genes in AKO-TAC hearts, while these pathways were enriched in WT-rMSA-TAC mice (Figure S4F and G, S5D and H). Untargeted lipidomics of cardiac tissues revealed higher total free fatty acids (FA) in AKO-TAC versus WT-TAC mice (Figure S10A), with distinct FA species alterations (Figure S10B). Levels of several triacylglycerol (TG) species were also elevated in AKO-TAC hearts (Figure S10C), along with sphingomyelins (SM), phosphatidylinositols (PI), and phosphatidylglycerols (PG) (Figure S10A and D–F). These lipid accumulations align with known mechanisms of HF progression involving lipotoxicity, mitochondrial dysfunction, and impaired FAO.^12^ Consistently, untargeted metabolomics further identified increased acylcarnitines (especially medium-to long-chain species) in AKO-TAC hearts, indicating impaired FAO flux (Figure S10G). Multi-omics integration suggested that albumin deficiency exacerbates metabolic inefficiency and lipid toxicity by disrupting FAO. At the molecular level, immunoblotting showed reduced PPARα protein levels in WT-TAC versus WT-Sham hearts, with further suppression in AKO-TAC hearts (Figure S11A). qRT-PCR confirmed downregulation of core FA metabolism genes in WT-TAC versus WT-Sham controls, including key transcription factor *Ppara*, FA uptake genes (*Slc27a1*, *Cd36*), FA activation and transport genes (*Acsl1*, *Cpt1b*, *Slc25a20*), and FAO enzymes (*Acadm*, *Acadl*, *Acadvl*, *Acaa2*) in WT-TAC versus WT-Sham hearts, with most further downregulated in AKO-TAC hearts (Figure 4A). *Cd36* expression was unchanged, indicating unaffected CD36-dependent FA uptake despite worsened FAO dysfunction (Figure 4A). Notably, *Slc27a1* (encoding FATP1) was significantly downregulated, potentially aggravating this metabolic defect. In addition, rMSA restored PPARα protein and FA metabolism gene expression in WT-rMSA-TAC versus WT-Saline-TAC hearts (Figure 4B and S11B). Similarly, AKO-rMSA-TAC mice exhibited elevated PPARα protein levels and alleviated FA metabolism gene suppression compared to AKO-Saline-TAC controls (Figure S11C and D). Notably, in AKO sham groups, rMSA also elevated expression of *Acsl1*, *Slc25a20*, and *Acadvl* (Figure S11D).

**Figure 4.**
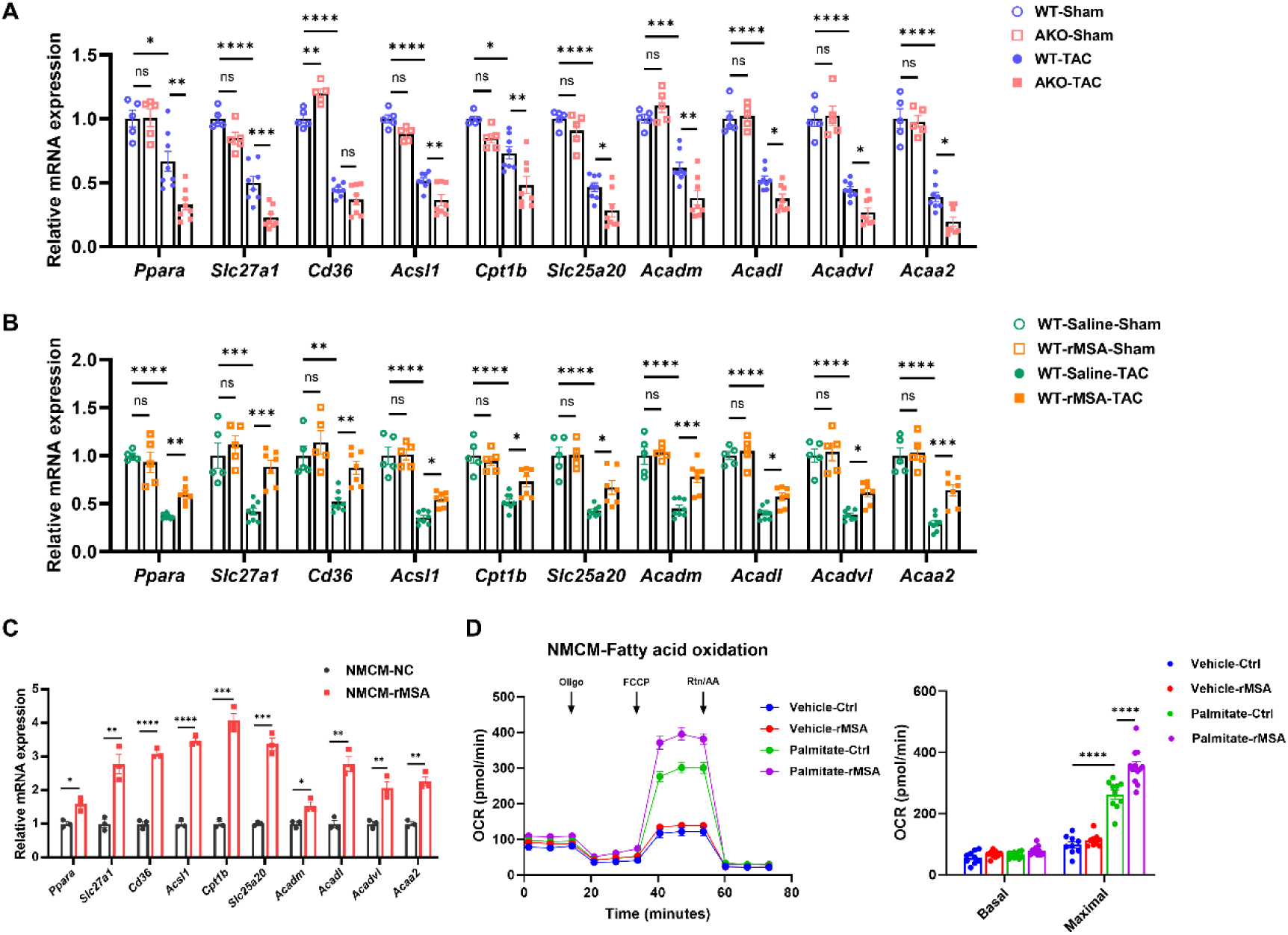
Albumin enhances cardiac fatty acid metabolism. (A) mRNA expression of cardiac fatty acid metabolism-associated genes from WT and AKO mice after sham or TAC 6 weeks (n = 5–8). (B) mRNA expression of cardiac fatty acid metabolism-associated genes from WT mice treated with rMSA or saline after sham or TAC 8 weeks (n = 5–8). (C) mRNA expression of fatty acid metabolism-associated genes in isolated primary neonatal mouse cardiomyocytes (NMCMs) treated with 5 mg/mL rMSA (NMCM-rMSA) or an equal volume of ddH₂O (NMCM-NC) (n = 3). (D) Measurement and quantification of fatty acid oxidation assay in NMCMs (n = 9– 12). Following a 24-hour serum starvation, NMCMs were treated with 5 mg/mL rMSA or an equal volume of ddH₂O (vehicle control) for 16 h. Four hours before the assay, cells were switched to a substrate-limited medium (low-glucose DMEM with 0.5 mM carnitine). One hour prior to measurement, the medium was replaced with fresh FAO assay medium (containing 2.5 mM glucose and 0.5 mM carnitine), and cells were incubated in a CO₂-free incubator at 37°C. The assay was initiated by adding a palmitate-BSA conjugate or BSA control (150 μM final BSA concentration). Sequential injections of oligomycin (2 μM), FCCP (3 μM), and antimycin A/rotenone (1 μM each) were performed. Basal and maximal respiration were calculated by subtracting the non-mitochondrial OCR. Fatty acid metabolism-associated genes including the key transcription factor (*Ppara*), fatty acid uptake-related genes (*Slc27a1* and *Cd36*), fatty acid activation/transport-related genes (*Acsl1*, *Cpt1b* and *Slc25a20*), and key enzymes involved in fatty acid β-oxidation (*Acadm*, *Acadl*, *Acadvl* and *Acaa2*) were detected in (A)–(C), normalized to *Actb* mRNA. Mean ± SEM, two-way ANOVA followed by Tukey’s multiple comparisons test (A, B and D) or unpaired two-tailed Student’s *t*-test (C). ns, not significant; \**p*<0.05, \*\**p*<0.01, \*\*\**p*<0.001, \*\*\*\**p*<0.0001.

Consistently, in isolated primary NMCMs, qRT-PCR confirmed the upregulation of key FA metabolism genes in rMSA-treated cells relative to controls (Figure 4C), and Seahorse FAO assays evaluating long-chain FAO capacity revealed that rMSA-treated cells exhibited higher maximal respiratory capacity than controls following exogenous palmitate supplementation (Figure 4D).

These data confirm that albumin preserves cardiac fatty acid metabolic homeostasis by maintaining transcriptional and functional FAO capacity, while its deficiency leads to toxic lipid accumulation and impaired bioenergetics in failing hearts.

### 2.6 rMSA attenuates phenylephrine (PE)-induced cardiomyocyte hypertrophy and mitochondrial dysfunction in NMCMs

To evaluate albumin’s direct cardiomyocyte effects, we used an *in vitro* PE-induced NMCM hypertrophy model. Immunoblotting showed rMSA attenuated PE-induced upregulation of hypertrophic and fibrotic markers (ANP, BNP, MYH7, Collagen I) (Figure 5A and B). Morphometric analysis confirmed that rMSA suppressed PE-induced cell surface area enlargement (Figure 5C and D). These findings validated albumin’s direct anti-hypertrophic effect on cardiomyocytes.

**Figure 5.**
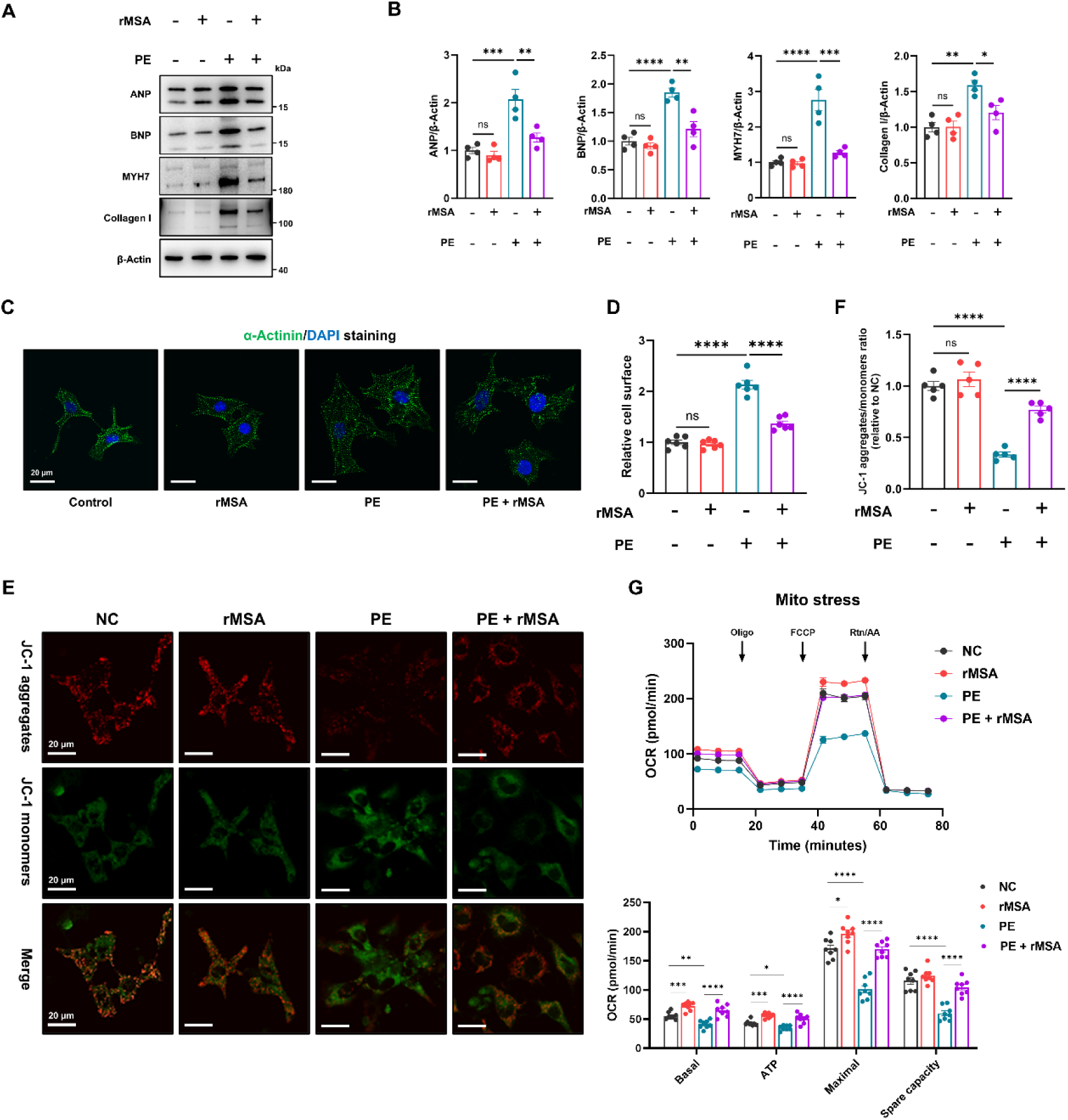
rMSA attenuates PE-induced cardiomyocyte hypertrophy and mitochondrial dysfunction in NMCMs. NMCMs were treated with 200 μM phenylephrine (PE) for 48 h under serum-free conditions to induce hypertrophy, with rMSA or an equal volume of ddH₂O (control). (A and B) Representative immunoblots (A) and quantification (B) of hypertrophy markers ANP, BNP, MYH7, and Collagen I protein levels in NMCMs, and β-Actin served as the loading control (n = 4). (C and D) Representative immunofluorescence staining of α-Actinin (C, scale bar, 20 μm) and statistical analysis of relative cardiomyocyte surface area (D, n = 6, ≥20 cells/dish) in NMCMs. (E and F) Representative JC-1 staining fluorescence images to assess mitochondrial membrane potential (E, scale bar, 20 μm) and the red (aggregates)/green (monomers) fluorescence intensity ratio of NMCMs (F, n = 5, ≥10 random fields/dish). Fluorescence images were captured by confocal microscopy and analyzed with ImageJ. (G) Mito stress test of NMCMs (n = 8). Mitochondrial respiratory parameters were assessed with sequential injections of oligomycin (2 μM), FCCP (1 μM), and antimycin A/rotenone (1 μM each). Basal respiration, ATP production, maximal respiration, and spare respiratory capacity were calculated by subtracting the non-mitochondrial OCR. Mean ± SEM, two-way ANOVA followed by Tukey’s multiple comparisons test. ns, not significant; \**p*<0.05, \*\**p*<0.01, \*\*\**p*<0.001, \*\*\*\**p*<0.0001.

The PE-induced model can effectively recapitulate mitochondrial dysfunction and metabolic dysregulation characteristic of HF.^33,34^ JC-1 staining revealed PE-induced mitochondrial membrane potential (ΔΨm) depolarization reflected by a reduced JC-1 aggregate/monomer ratio, and rMSA treatment prevented this ΔΨm loss (Figure 5E and F). Furthermore, the Seahorse mitochondrial stress test showed that rMSA significantly enhanced basal respiration, ATP production, maximal respiration, and spare respiratory capacity in PE-induced NMCMs, with mild baseline improvements (Figure 5G).

These data confirm that albumin directly protects cardiomyocytes by attenuating pathological hypertrophy and preserving mitochondrial function *in vitro*, underscoring its therapeutic potential for pathological cardiac remodeling.

### 2.7 Albumin activates the PKA-LKB1-AMPK signaling axis

To explore the underlying mechanisms, transcriptomic KEGG analysis revealed significant enrichment of the AMPK signaling pathway in key comparisons: genes downregulated in AKO-TAC versus WT-TAC hearts, and genes upregulated in WT-rMSA-TAC versus WT-Saline-TAC hearts (Figure S4E and S5C). Heatmaps further highlighted consistent AMPK-related expression patterns (Figure S12), implicating this pathway in albumin-mediated cardioprotection and mitochondrial homeostasis regulation.

AMP-activated protein kinase (AMPK) is a cardiac energy sensor and regulator of mitochondrial function and substrate metabolism.^35,36^ To investigate the effects of rMSA on the AMPK signaling pathway, serum-starved NMCMs were treated with rMSA. Immunoblotting showed that rMSA treatment time-and dose-dependently activated phosphorylation of AMPK and its downstream target ACC, and elevated PGC-1α expression (Figure 6A and B). This activation was abolished by the AMPK inhibitor Compound C (CpC) (Figure 6C). AMPK activation can occur via multiple upstream kinases, including LKB1 and CAMKK2.^35^ Immunoblotting showed that rMSA treatment increased phosphorylation of LKB1 at Ser431 (corresponding to human Ser428) in a time- and dose-dependent manner, but did not affect CAMKK2 expression (Figure 6D and E). siRNA-mediated knockdown of LKB1 blocked rMSA-induced AMPK and ACC phosphorylation, as well as PGC-1α upregulation (Figure 6F), confirming LKB1 as the critical upstream kinase. As LKB1 Ser431 phosphorylation can be mediated by PKA or PKCζ,^37,38^ we sought to identify the kinase responsible for rMSA-induced LKB1-AMPK activation. rMSA treatment increased PKA substrate phosphorylation in a time- and dose-dependent manner (peaking at 5 mg/mL, then declining), while PKCζ Thr410 phosphorylation remained unchanged (Figure 6G, H and S13). Furthermore, inhibition of PKA with H89 blocked rMSA-induced phosphorylation of LKB1, AMPK, and ACC, as well as PGC-1α expression (Figure 6I), establishing PKA as the upstream activator. Transcriptomic KEGG analysis also revealed significant enrichment of the cAMP signaling pathway in two comparisons: genes downregulated in AKO-TAC versus WT-TAC hearts, genes upregulated in WT-rMSA-TAC versus WT-Saline-TAC hearts (Figure S4E and S5C). Heatmaps further highlighted the cAMP-related expression patterns (Figure S14). *In vitro* experiments using rMSA at varying concentrations and durations, combined with inhibitor and siRNA knockdown approaches, demonstrate that rMSA enhances mitochondrial homeostasis by activating the PKA-LKB1-AMPK signaling axis in cardiomyocytes.

**Figure 6.**
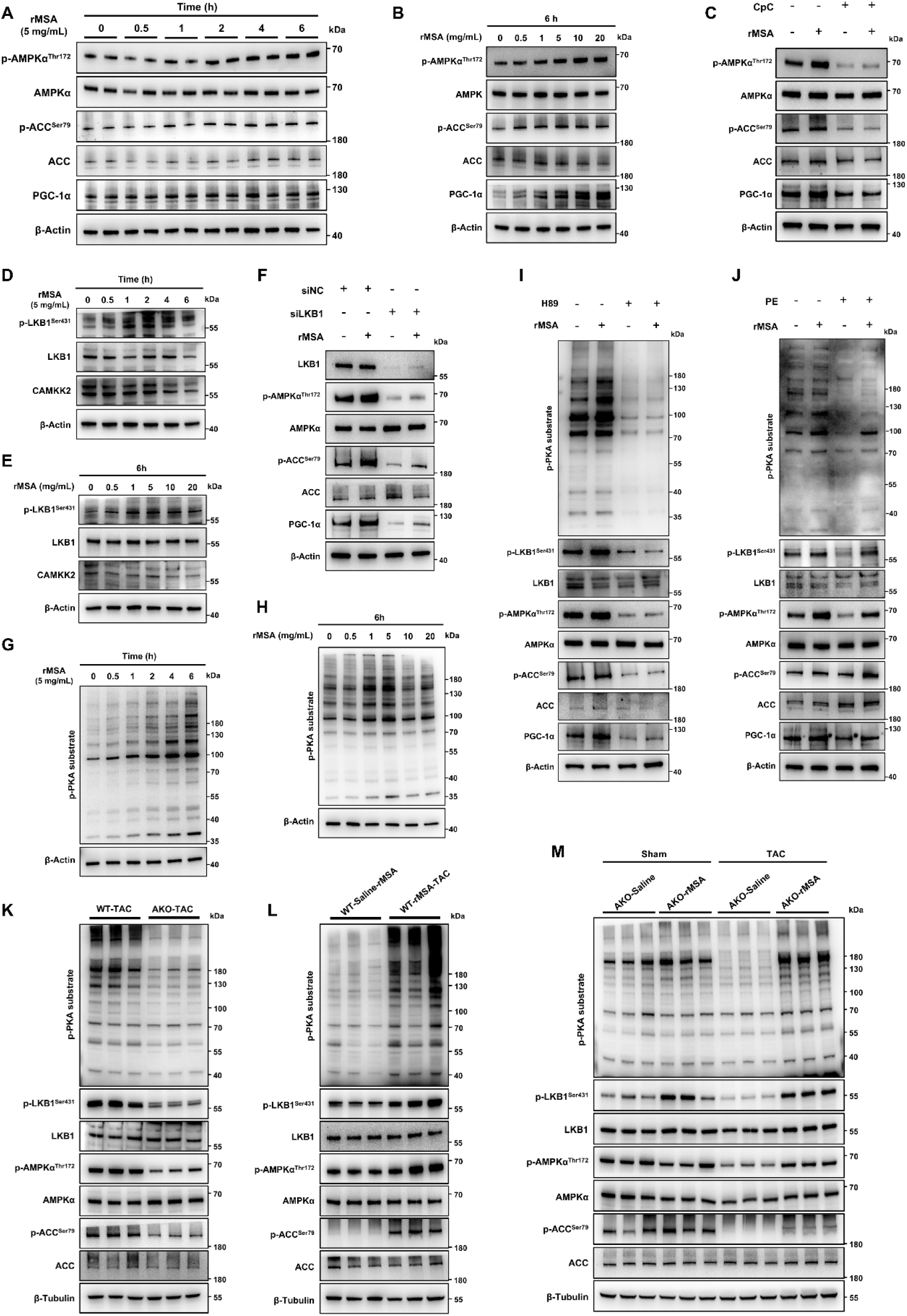
Albumin activates the PKA-LKB1-AMPK signaling axis in NMCMs. (A and B) Representative immunoblots of phospho-/total AMPK and ACC, and PGC-1α in NMCMs treated with rMSA over a time course (0, 0.5, 1, 2, 4 and 6 h) (A), and a concentration gradient (0, 0.5, 1, 5, 10, 20 mg/mL) for 6 h (B). (C) Representative immunoblots of phospho-/total AMPK and ACC, and PGC-1α, in NMCMs treated with 5 mg/mL rMSA for 6 h, with or without 5 μM Compound C (CpC, AMPK inhibitor) (n = 4). (D and E) Representative immunoblots of phospho-/total LKB1 and CAMKK2 in NMCMs treated with rMSA over a time course (D), and a concentration gradient for 6 h (E). (F) Representative immunoblots of LKB1, phospho-/total AMPK and ACC, and PGC-1α in NMCMs transfected with siLKB1 or siNC for 48 h, followed by 5 mg/mL rMSA for 6 h (n = 4). (G and H) Representative immunoblots of PKA substrate phosphorylation in NMCMs treated with rMSA over a time course (G), and a concentration gradient for 6 h (H). (I) Representative immunoblots of PKA substrate phosphorylation, phospho-/total LKB1, AMPK, ACC, and PGC-1α in NMCMs treated with 5 mg/mL rMSA for 6 h, with or without 5 μM H89 (PKA inhibitor) (n = 4). (J) Activation of the PKA-LKB1-AMPK signaling axis was assessed by representative immunoblots of phosphorylated PKA substrate, phospho-/total LKB1, AMPK, ACC, and PGC-1α protein levels in the PE-induced cardiomyocyte hypertrophy model (n = 4). Quantification is shown in Figure S15. (K–M) Activation of the PKA-LKB1-AMPK signaling axis was assessed by representative immunoblots and quantification of phosphorylated PKA substrate, phospho-/total LKB1, AMPK, ACC, and PGC-1α protein levels in cardiac tissues from WT and AKO mice after TAC 6 weeks (K, n = 4), from WT mice treated with rMSA or saline after TAC 8 weeks (L, n = 4), from AKO mice treated with rMSA or saline after sham or TAC 8 weeks (M, n = 4). Quantification is shown in Figure S15. (A–J) β-Actin and (K–M) β-Tubulin served as the loading controls.

Thereafter, we verified this signaling axis in PE-induced hypertrophic NMCMs. rMSA restored suppressed phosphorylation of PKA substrates, LKB1, AMPK, and ACC, as well as downregulation of PGC-1α expression (Figure 6J and S15A). We next validated this signaling axis in TAC-induced mice with distinct genotypes and treatment regimens established in this study. AKO-TAC hearts showed reduced phosphorylation of PKA substrates, LKB1, AMPK, and ACC compared to WT-TAC controls (Figure 6K and S15B). Conversely, rMSA treatment elevated these phosphorylation levels in WT-rMSA-TAC versus WT-Saline-TAC hearts (Figure 6L and S15C). Similarly, AKO-rMSA-TAC mice exhibited elevated phosphorylation levels of these proteins compared to AKO-Saline-TAC controls (Figure 6M and S15D). Notably, rMSA also increased phosphorylation of PKA substrates and LKB1 in AKO-Sham mice but did not alter AMPK and ACC phosphorylation, likely due to homeostatic regulation in the unstressed state (Figure 6M and S15D). These data indicate that albumin can activate the PKA-LKB1-AMPK axis under pressure overload, suggesting its therapeutic potential for HF.

To determine whether the PKA-LKB1-AMPK axis mediates the cardioprotective effects of albumin against PE-induced cardiomyocyte hypertrophy, we conducted AMPK or PKA inhibition and LKB1 knockdown experiments in NMCMs. AMPK inhibition with CpC significantly abrogated the protective effect of rMSA against PE-induced hypertrophy (Figure 7A). Similarly, genetic knockdown of LKB1 impaired rMSA’s ability to reduce hypertrophic marker expression (Figure 7B). Furthermore, PKA inhibition with H89 eliminated rMSA’s protective effects (Figure 7C). Collectively, these data demonstrate that albumin can mitigate PE-induced NMCM hypertrophy via the PKA-LKB1-AMPK axis.

**Figure 7.**
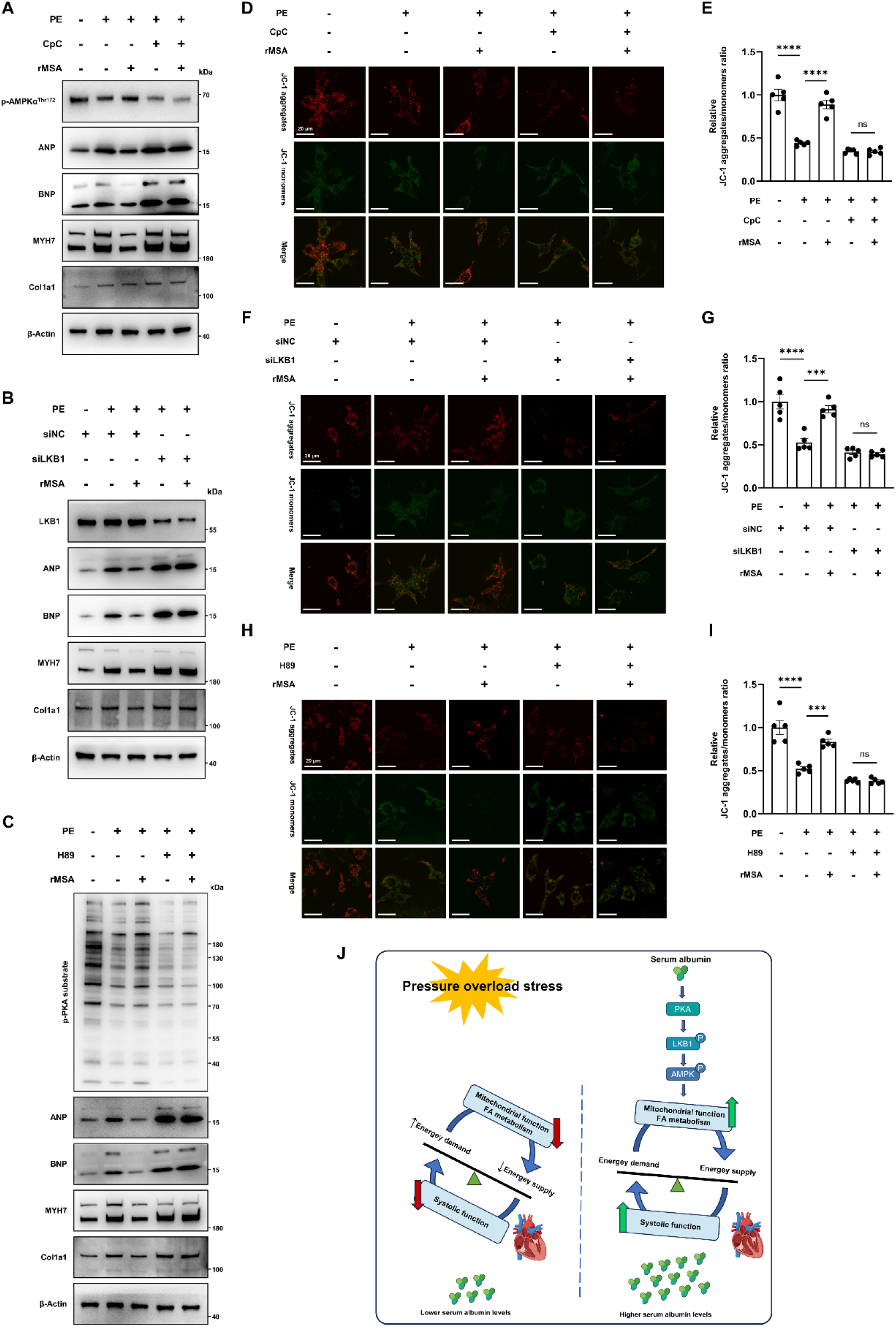
The PKA-LKB1-AMPK signaling axis mediates rMSA protection against PE-induced cardiomyocyte hypertrophy via maintaining mitochondrial homeostasis. (A–C) Representative immunoblots of hypertrophy markers (ANP, BNP, MYH7, and Collagen I) in PE-induced NMCMs after treatment with rMSA in the presence or absence of CpC (A, n = 4), after transfection with siLKB1 or siNC for 48 h (B, n = 4), and after treatment with rMSA in the presence or absence of H89 (C, n = 4). β-Actin served as the loading control. (D–I) Representative JC-1 staining fluorescence images (scale bar, 20 μm) to assess mitochondrial membrane potential and the red (aggregates)/green (monomers) fluorescence intensity ratio in PE-induced NMCMs after treatment with rMSA in the presence or absence of CpC (D and E, n = 5), after transfection with siLKB1 or siNC for 48 h (F and G, n = 5), and after treatment with rMSA in the presence or absence of H89 (H and I, n = 5), with ≥10 random fields analyzed per dish. Fluorescence images were captured by confocal microscopy and analyzed with ImageJ. (E, G and I) Mean ± SEM, one-way ANOVA followed by Tukey’s multiple comparisons test. ns, not significant; \*\*\**p*<0.001, \*\*\*\**p*<0.0001. (J) A schematic illustration. Reduced serum albumin levels accelerate heart failure progression. Conversely, albumin activates the PKA-LKB1-AMPK signaling axis, which enhances mitochondrial function and fatty acid metabolism. This improvement in mitochondrial homeostasis ultimately alleviates pressure overload-induced heart failure.

To further explore whether this axis mediates the regulatory effects of rMSA on mitochondrial homeostasis during PE-induced cardiomyocyte hypertrophy, we assessed mitochondrial membrane potential (ΔΨm) via JC-1 staining following AMPK inhibition with CpC, siRNA-mediated LKB1 knockdown, or PKA inhibition with H89. Notably, blockade of AMPK activity, genetic ablation of LKB1, or suppression of PKA signaling each markedly reduced ΔΨm and aggravated mitochondrial dysfunction under PE challenge, and rMSA failed to restore the impaired ΔΨm upon disruption of the PKA-LKB1-AMPK cascade (Figure 7D–I). These results indicate that albumin preserves ΔΨm and maintains mitochondrial homeostasis against PE-induced mitochondrial dysfunction in a PKA-LKB1-AMPK axis-dependent manner.

Collectively, this study demonstrates that albumin can activate the PKA-LKB1-AMPK signaling axis, enhancing mitochondrial function and fatty acid metabolism, thereby improving mitochondrial homeostasis and alleviating pressure overload-induced HF (Figure 7J).

## 3. Discussion

In this study, we identified a critical cardioprotective role of serum albumin in pressure overload-induced HF. The established clinical link between low serum albumin levels and poor HF prognosis prompted us to investigate the direct role of albumin. In our mouse model, TAC surgery similarly reduced serum albumin levels and worsened cardiac dysfunction and remodeling in WT mice. Direct evidence was obtained in AKO mice, which developed more severe HF progression after TAC. Crucially, rMSA supplementation restored circulating albumin, ameliorated injury in TAC-induced WT mice, and attenuated disease progression in TAC-induced AKO mice, confirming its therapeutic potential. rMSA also directly suppressed cardiomyocyte hypertrophy induced by PE *in vitro*. By integrating *in vivo* and *in vitro* models, we found that albumin could activate the PKA-LKB1-AMPK signaling axis. This activation orchestrated a comprehensive improvement in mitochondrial homeostasis, enhancing respiratory function, quality control, and fatty acid oxidation. Thus, albumin effectively corrected the key cardiac energy imbalance underlying disease progression. Our findings uncover a novel mechanism by which albumin preserves cardiac function, establishing recombinant albumin as a viable therapeutic approach for HF.

To explore the role of albumin, we utilized AKO mice, a model exhibiting complete absence of serum albumin that serves as an extreme manifestation of hypoalbuminemia. Several studies demonstrated that young AKO mice exhibit a slight but significant decrease in total serum protein and non-esterified fatty acids, along with elevated total bilirubin, lipase, calcium, ALT, total cholesterol, HDL, LDL, AST, iron, and triglycerides, and this pattern reflects systemic hyperlipidemia and resembles human analbuminemia.^39-41^ Our previous work further revealed that AKO mice develop spontaneous obesity and progressive non-alcoholic fatty liver disease (NAFLD) with aging, accompanied by marked dysregulation of hepatic lipid metabolism.^42^ However, we observed no significant impairment in baseline cardiac systolic function or elevation of NT-proBNP levels in the absence of pathological stress in this study, regardless of age. We hypothesize that upregulated blood proteins compensate for albumin’s basal functions in AKO mice but are insufficient under pathological stress. Given that obesity and dyslipidemia are established HFpEF risk factors, further investigation of diastolic dysfunction in this model is warranted.^43^

Our study reveals that albumin can positively regulate MQC mechanisms. While excessive mitochondrial fission is commonly linked to HF progression,^44^ our TAC model showed a generalized suppression of MQC, including reduced biogenesis, fusion, fission, and mitophagy, leading to impaired turnover and accumulation of damaged mitochondria. These findings contrasted with reports emphasizing hyperactive fission. We propose that pathological stress may suppress fission as an adaptive response to limit mitochondrial fragmentation; however, dysfunction of other MQC pathways ultimately compromises this adaptation. Additionally, fission is essential for segregating and clearing damaged mitochondria via mitophagy, and its excessive suppression can be detrimental.^11^ Thus, precisely balancing MQC regulation is essential for potential clinical translation. Furthermore, our data show that albumin upregulates PGC-1α, which promotes mitochondrial biogenesis and serves as a PPARα co-activator to enhance fatty acid oxidation. This coordinated enhancement of mitochondrial capacity and substrate utilization establishes a mutually reinforcing cycle that supports cardiac energy homeostasis under stress.

Accumulating evidence has long indicated the intrinsic association between albumin and mitochondrial homeostasis across diverse species and cell types. An early 1958 study showed that serum albumin is essential for oxidative phosphorylation in insect mitochondria.^45^ Similarly, bovine serum albumin reportedly enhances substrate oxidation, particularly succinate metabolism, in mammalian brain mitochondria.^46^ Moreover, recent work validated that albumin supplementation restores normal mitochondrial morphology in damaged endothelial cells from cirrhotic patients.^47^ Despite decades of evidence that albumin regulates mitochondrial function, its mechanism in cardiomyocytes remains unclear. Here, we identify the PKA-LKB1-AMPK axis as a previously unrecognized pathway by which albumin preserves mitochondrial homeostasis and alleviates cardiomyocyte hypertrophy.

The activation of AMPK enhances metabolic flexibility by regulating these processes, thereby attenuating pathological cardiac hypertrophy and HF progression, thus providing a molecular basis for metabolic interventions in cardiovascular diseases.^48-50^ AMPK is phosphorylated and activated mainly by LKB1 under energy stress, forming the LKB1-AMPK axis that is crucial for maintaining cardiac energy homeostasis.^35,51^ Further upstream, PKA, a cAMP-dependent kinase, can phosphorylate LKB1 at Ser428, thereby integrating extracellular signals with intracellular responses to modulate metabolism, gene expression, and cell survival.^52^ Impaired PKA signaling, including its role in regulating cardiac contractility, is a hallmark of HF, highlighting the critical importance of this regulatory cascade.^52,53^ The PKA-LKB1-AMPK axis plays a critical role in cardiac hypertrophy or other pathophysiology.^54,55^ While our data indicate that albumin can activate this pathway to ameliorate mitochondrial dysfunction in HF, the precise mechanism initiating PKA activation remains unclear, as no studies have elucidated how albumin activates the cAMP-PKA pathway. In cardiomyocytes, cAMP-PKA signaling is classically triggered through Gαs-coupled adenylate cyclases (ACs), with cAMP levels balanced by the activity of phosphodiesterases (PDEs).^53^ We hypothesize that albumin modulates cAMP by directly activating ACs or inhibiting PDEs. In addition, albumin’s canonical antioxidant and anti-inflammatory effects may indirectly maintain PKA function. A further possibility involves the biological activity of molecules carried by albumin. Notably, downregulated β2-adrenergic receptor (β2AR) expression and impaired membrane localization reduce cAMP production, suppressing PKA activity and worsening cardiovascular dysfunction.^56^ β2AR internalization and disrupted cAMP-PKA signaling are associated with caveolin-3 (CAV3) dysregulation, which normally anchors β2AR in caveolae to maintain membrane stability.^57-59^ Our previous work found that albumin promotes caveolin-1 (CAV1)-dependent caveolae formation in hepatocytes, enhancing receptor-mediated signaling.^20^ Given the structural homology between CAV1 and CAV3, we propose that albumin may similarly stabilize β₂AR localization in cardiomyocytes via CAV3. This mechanism could attenuate β₂AR internalization, restore adrenergic responsiveness, and facilitate downstream LKB1-AMPK activation. Although cardiomyocytes lack canonical albumin receptors (e.g., gp60, FcRn), CAV3 may mediate albumin uptake through non-canonical endocytic pathways.^60^

Our findings establish albumin as a key regulator of mitochondrial homeostasis in HF. While this study focuses on its direct effects on cardiomyocytes, HF pathophysiology involves complex intercellular crosstalk among fibroblasts, immune cells, and endothelial cells. Emerging evidence indicates that albumin exerts cardioprotection via multi-target mechanisms, such as protecting vascular endothelial cells and maintaining coronary endothelial glycocalyx integrity.^61-63^ To clarify albumin’s therapeutic potential in HF, future studies should use multicellular models and investigate intercellular signaling pathways to refine treatment strategies.

HF patients often exhibit hypoalbuminemia, linked to adverse outcomes due to altered vascular permeability, increased catabolism, and reduced synthesis.^23^ Intravenous albumin is commonly used for perioperative fluid resuscitation in cardiac surgery to address hemodynamic instability by alleviating edema and serous effusions through osmotic regulation, hemodynamic modulation, and enhanced diuresis.^64^ While albumin infusion has been attempted for HF management, its clinical efficacy remains inconsistent, and its use is controversial due to risks of volume overload, renal dysfunction, and allergic reactions.^65-67^ This is particularly relevant in advanced HF, where impaired cardiac compensatory capacity limits albumin tolerance, with risks outweighing its potential benefits. In contrast, our study demonstrated that early albumin administration might delay HF progression by improving mitochondrial function and energy metabolism. Furthermore, while blood-derived albumin carries risks of contamination and modification, recombinant serum albumin presents a promising alternative.^68^ We thus investigated whether recombinant albumin is functionally superior to plasma-derived albumin. rMSA activated the PKA-LKB1-AMPK axis more effectively than plasma-derived mouse serum albumin (pMSA) in NMCMs, evidenced by higher phosphorylation of PKA substrates, LKB1, AMPK and ACC and greater PGC-1α expression (Figure S16A). Consistently, rMSA better attenuated PE-induced hypertrophy, as shown by more pronounced reductions in hypertrophic marker protein levels compared to pMSA (Figure S16B). These data demonstrate that rMSA can more effectively activate cardioprotective signaling and mitigate pathological hypertrophy in NMCMs. Future research should focus on optimizing albumin quality and dosing regimens, particularly in specific patient subgroups, such as those with hypoalbuminemia and mitochondrial dysfunction.

HSA, a multifunctional plasma protein, undergoes multiple modifications closely linked to systemic oxidative stress and disease progression. Notably, the redox state of HSA, particularly the oxidation of its Cys34 thiol group, a key antioxidant component, serves as a critical biomarker for oxidative stress severity and cardiovascular mortality in conditions such as hemodialysis and acute lung injury.^69,70^ In recent years, the concept of "effective albumin concentration" has emerged, highlighting that its decline reflects not only quantitative loss but also qualitative dysfunction from various modifications, underscoring the need for quality-focused assessment.^71^ Our previous work demonstrated that murine aging is associated with progressive albumin modifications, characterized by decreased free thiol levels and increased carbonylation, advanced glycation end-products and homocysteinylation, which collectively impair its physiological function.^19^ Our group defines the concept of "young and undamaged" for proteins to evaluate their quality based on these four parameters, and proposes that recombinant serum albumin is superior to plasma-derived albumin in quality.^19^ Moreover, the recombinant human albumin (rHSA) developed by our collaborators offers high purity, low immunogenicity, and excellent batch consistency, with these properties providing a solid foundation for its clinical translation.

This study supports expanded applications of recombinant albumin in cardiovascular diseases, particularly HF, by elucidating its role in mitochondrial homeostasis and cardioprotection, while providing translational insights for novel therapy development.

### Translational Perspective

Heart failure (HF) remains a major clinical challenge with limited therapeutic options to alleviate disease progression. Based on the established correlation between low serum albumin levels and adverse HF outcomes, this study provides evidence that albumin depletion directly aggravates HF progression rather than merely serving as a disease biomarker. We show that recombinant albumin supplementation activates the PKA-LKB1-AMPK signaling axis for cardioprotection, restoring mitochondrial function and energy metabolism in failing hearts. These findings position recombinant serum albumin as a promising therapeutic strategy, particularly for HF patients with hypoalbuminemia, offering a novel approach to targeting cardiac energy imbalance.

## 4. Methods

### Experimental animals

The wild type (WT, C57BL/6JNifdc) mice used in this study were purchased from Charles River, and the albumin knockout (AKO, C57BL/6J-*Alb^em8Mvw^*/MvwJ, Strain #: 025200) mice strain was purchased from The Jackson Laboratory (JAX). Rearing of all mice and breeding of AKO mice were performed at the Laboratory Animal Resources Center of Tsinghua University. Following professional transport, mice were housed in Tsinghua University’s specific pathogen free (SPF)-level animal facility with 12-h light/dark cycles at 20-24°C and had free access to food and drinking water. All procedures complied with Tsinghua University Institutional Animal Care and Use Committee (IACUC) guidelines. Male mice received standard diet (SWC9102, Xietong Shengwu) and sterile water. Bedding was changed weekly with daily health monitoring. Acclimated animals were stabilized for 1 week pre-experiment. Any health-compromised mice were excluded from survival studies. AKO mice were generated by crossing homozygous *Alb*^-/-^males with heterozygous *Alb*^+/-^ females following the JAX-recommended breeding strategy. Mouse tails were collected at 3 weeks postnatally for genotyping using Quick Genotyping Assay Kit for Mouse Tail (D7283S, Beyotime) and Sanger sequencing with JAX-recommended primers: forward 5’-TTCCAAACCTCCGTGAAAAC-3’, reverse 5’-ATTCCCAGACTTGGTGTTGG-3’. AKO males were selectively utilized for experiments after genotype confirmation. In this study, male WT and AKO mice aged 2 months or 24 months underwent echocardiography followed by blood and cardiac tissue collection. Additional male WT and AKO (10–12 weeks, 25–30 g) were subjected to transverse aortic constriction (TAC) to establish pressure-overload heart failure models followed by echocardiography, and blood and cardiac tissue collection.

After echocardiography, mice were anesthetized with intraperitoneal avertin (20 μL/g), followed by orbital blood collection. Blood samples were coagulated at room temperature for 45 min, centrifuged (3000 rpm, 15 min, 4°C), and the serum supernatants stored at -80°C. Hearts were perfused with PBS via peristaltic pump following thoracotomy, then excised, cleaned, and weighed. Heart weight was normalized to tibia length. Specimens were photographed on calibrated plates. Left ventricular tissues were partitioned for: 1) fixation in 4% paraformaldehyde for 24 h (for histopathology); 2) preservation in RNA stabilization buffer (MF150, Mei5bio) (for transcript analysis); 3) fixation in glutaraldehyde (for transmission electron microscopy); or 4) flash-freezing at -80°C (for immunoblotting, lipidomics, metabolomics, etc.). Additionally, primary neonatal mouse cardiomyocytes (NMCMs) were isolated from 1-3-day-old WT pups, irrespective of sex.

### Transverse aortic constriction (TAC) surgery

Mice were anesthetized via isoflurane inhalation using a rodent ventilator, with anesthesia depth confirmed by loss of corneal reflex, muscle relaxation, and flaccid paralysis, and then secured supine on a surgical board. Following tracheal intubation connected to mechanical ventilation (105–110 breaths/min, tidal volume 2–3 mL), the surgical site was prepared by removing fur from the neck to the second and third rib region with depilatory cream and disinfecting with alcohol swabs. A left parasternal incision exposed the second rib, which was resected to access the ascending aorta. Aortic constriction was induced by ligating the aorta between the first and second aortic arch branches with a 6-0 silk suture alongside a customized steel cylinder (0.4 mm in diameter); the cylinder was removed post-ligation to create circumferential constriction, followed by layered closure with 4-0 sutures. Postoperative analgesia and thermal support were provided, with continuous monitoring during recovery. Sham-operated mice underwent identical anesthesia, thoracotomy, and suturing without aortic constriction. This standardized protocol ensured consistent pressure overload modeling while minimizing perioperative complications.

### Echocardiographic analyses

A high-frequency, high-resolution digital imaging system (VisualSonics Vevo2100, Fujifilm) was used for echocardiography. Following respiratory anesthesia, mice were positioned supine on the ultrasound platform with chest and upper abdominal fur removed using depilatory cream. The paws were secured to a conductive gel-coated stage for stable imaging. A parasternal long-axis view was optimized by angling the probe toward the left chest to visualize the left atrium and ventricular inflow tract, ensuring maximal left ventricular (LV) dimensions. B-mode imaging was performed to guide M-mode acquisition at the level of the mitral valve chordae. Key parameters, including left ventricular ejection fraction (LVEF), left ventricular fractional shortening (LVFS), left ventricular end-diastolic volume (LV Vol;d), left ventricular end-systolic volume (LV Vol;s), left ventricular internal diameter at end-diastole (LVID;d), left ventricular internal diameter at end-systole (LVID;s), left ventricular posterior wall thickness at end-diastole (LVPW;d), left ventricular posterior wall thickness at end-systole (LVPW;s), left ventricular anterior wall thickness at end-diastole (LVAW;d), left ventricular anterior wall thickness at end-systole (LVAW;s), left ventricular mass (LV mass), and corrected left ventricular mass (LV mass cor), were calculated from ≥5 consecutive cardiac cycles and averaged. All procedures were conducted under standardized respiratory settings to ensure reproducibility.

### Administration of recombinant mouse serum albumin (rMSA)

In this study, young, undamaged, and ultrapure rMSA (Shenzhen Protgen, Ltd.) was administered via tail vein injection (*i.v.*) to mice at a dose of 1.5 mg per gram of body weight (1.5 mg/g), with an equal volume of saline used as the vehicle control. A staged dosing regimen was implemented: 1) preoperative phase: mice were administered rMSA twice (48 hours apart) starting 4 days before TAC to elevate baseline albumin levels. 2) postoperative phase: dosing was continued with additional injections every 3 weeks for two cycles to sustain and further elevate albumin levels. The identical rMSA dosing regimen was applied to AKO mice to evaluate albumin restoration effects. All injections were performed uniformly by a single operator to ensure consistency.

### N-terminal pro-brain natriuretic peptide (NT-proBNP) ELISA

Serum NT-proBNP concentrations were measured using a mouse NT-proBNP ELISA Kit (E-EL-M0834, Elabscience). Briefly, reagents were equilibrated at room temperature, and a standard curve was prepared via serial dilutions. Standards, blanks, and diluted serum samples were loaded onto a pre-coated plate and incubated at 37°C for 90 min. After incubation with biotinylated antibody and HRP conjugate, the plate was washed, and TMB substrate was added for color development. The reaction was stopped, and optical density was measured at 450 nm using a microplate reader.

### Tissue paraffin embedding and sectioning

Tissue samples were fixed in 4% paraformaldehyde for 24 h at room temperature, rinsed with PBS, and dehydrated through a graded ethanol series. Tissues were cleared in xylene and infiltrated with molten paraffin at 60°C. Paraffin-embedded tissues were sectioned into 5 μm slices using a microtome, floated on a 40°C water bath, and mounted on glass slides for drying.

### Hematoxylin and eosin (H&E) staining

An H&E Staining Kit (C0105S, Beyotime) was used. Sections were dewaxed, rehydrated through a graded ethanol series, and stained with hematoxylin for 5 min, followed by eosin counterstaining. After dehydration and clearing in xylene, slides were mounted with neutral resin and scanned for histological analysis.

### Masson’s trichrome staining

Following dewaxing and rehydration, sections were stained using a Masson’s Trichrome Staining Kit (C0189S, Beyotime) with hematoxylin for 5 min, Ponceau-acid fuchsin for 10 min, and aniline blue for 1 min. Collagen fibers (stained blue) were quantified using ImageJ software to determine the percentage of fibrosis area.

### Sirius red staining

Sections were dewaxed, rehydrated, and stained with PicroSirius Red (DC0041, Leagene) for 30 min, followed by hematoxylin counterstaining. Collagen fibers (stained red) were quantified using ImageJ software to assess collagen volume fraction.

### Wheat germ agglutinin (WGA) staining

After dewaxing and rehydration, antigen retrieval was performed using citrate buffer (P0081, Beyotime). Sections were then incubated with WGA-FITC (L4895, Sigma) working solution for 1 h in the dark. Sections were imaged using a confocal laser scanning microscope (FV3000, Olympus). ImageJ software was applied to quantify the cross-sectional area of individual cardiomyocytes. Three discontinuous tissue sections per mouse were analyzed, with a minimum of 200 fibers measured to calculate the mean cross-sectional area.

### Transmission electron microscopy (TEM) analysis of mitochondrial ultrastructure

Mitochondrial ultrastructure in mouse cardiac tissues was analyzed by TEM (HT7800, Hitachi). Tissue blocks (1 mm³) were immediately fixed in 2.5% glutaraldehyde at 4°C for 2 h, followed by four washes with 0.1 M phosphate buffer (pH 7.4). Post-fixation was performed in 1% osmium tetroxide containing 1.5% potassium ferrocyanide for 30 min, with subsequent ddH₂O rinses. Samples were stained with 1% uranyl acetate overnight, dehydrated through graded ethanol series (50%, 70%, 80%, 100%), and embedded in epoxy resin after three 100% ethanol washes. Polymerization was completed at 60°C for 24 h. Ultrathin sections (70 nm) were cut with a diamond knife, mounted on Formvar-coated copper grids, and examined under TEM at 80 kV acceleration voltage.

### Quantitative real-time PCR (qRT-PCR)

Total RNA was extracted from cardiac tissues using the HiPure Fibrous RNA Plus Kit (R4124, Magen) and from liver tissues and cells using the HiPure Universal RNA Mini Kit (R4130, Magen), following the manufacturers’ protocols. RNA was purified using mini columns, washed, and eluted in RNase-free water. Concentration and purity were measured using a NanoDrop spectrophotometer, and RNA samples were stored at -80°C for subsequent analysis. First-strand cDNA synthesis was performed using the HiScript III 1st Strand cDNA Synthesis Kit (R312-01, Vazyme). For qRT-PCR, cDNA was diluted 10-fold with ddH_2_O, and reactions were prepared in triplicate using Taq Pro Universal SYBR qPCR Master Mix (Q712-02, Vazyme). Amplification was conducted using LightCycler 480 Instrument II (Roche). The *Actb* gene was used as the endogenous control. Relative gene expression levels were determined using the 2^−ΔΔCt method. Primer sequences used for qRT-PCR in this study are listed in Table S6.

### Mitochondrial DNA Copy Number Analysis

Approximately 20 mg of cardiac tissues was processed for the analysis. Total DNA, including mitochondrial DNA (mtDNA) and nuclear DNA (nDNA), was extracted using the TIANamp Genomic DNA Kit (DP304-03, TIANGEN). The copy numbers of mtDNA (targeting *mtND1*) and nDNA (targeting *HK2*) were quantified by qPCR on a LightCycler 480 II (Roche) using Taq Pro Universal SYBR qPCR Master Mix. Primer sequences were provided in Table S6.

### Western Blotting

For tissue samples, 20–30 mg of liquid nitrogen-flash-frozen tissue (stored at -80°C) from each mouse was homogenized on ice with complete RIPA buffer (P0038, Beyotime) containing protease and phosphatase inhibitors (P1048, Beyotime) using an electric homogenizer (3–4 pulses, ≤5 s each). After incubation on ice for 10 min, lysates were centrifuged (12,000 × *g*, 5 min, 4°C), and the supernatants were collected. For cell lysis, cultures were washed with ice-cold PBS, lysed with RIPA buffer (prepared as above), incubated on ice for 10 min, gently scraped, centrifuged under identical conditions, and the supernatants were collected. Protein concentrations were determined using the Pierce™ BCA Protein Assay Kit (23225, Thermo Scientific). A standard curve was prepared using BSA solutions (62.5–2000 μg/mL). Samples and standards were mixed with the BCA working reagent, incubated at 37°C for 30 min, and the absorbance at 562 nm was measured using a microplate reader for concentration calculation. For cardiac tissues and cell samples, protein concentrations were normalized using lysis buffer based on BCA assay results, mixed with 5× SDS-PAGE loading buffer (P0015L, Beyotime) and denatured at 95°C for 10 min. Serum samples were diluted 400-fold and treated similarly. Samples were either loaded directly onto SDS-PAGE gels or stored at -80°C. Electrophoresis was performed using 4–12% polyacrylamide gels (P41215, Lablead) at 120 V for 60 min. For protein transfer, activated PVDF membranes (IPVH00010, Millipore) and gels were assembled into a transfer sandwich and subjected to wet transfer at 90 V for 60–90 min (duration adjusted based on target protein molecular weight). Membranes were blocked with 5% skim milk (P0216, Beyotime) in TBST (G0001, Servicebio) for 60 min at room temperature, followed by incubation with primary antibodies overnight at 4°C. After three 10-min washes with TBST, membranes were incubated with HRP-conjugated secondary antibodies for 60 min at room temperature. Following additional washes, target proteins were visualized using chemiluminescent substrate either SuperSignal West Femto (34095, Thermo Scientific) or Western Blotting Luminol Reagent (sc-2048, Santa Cruz) and imaged with a chemiluminescence detection system. Band intensities were quantified using ImageJ software. Antibodies used in this study are listed in Table S7.

### RNA sequencing and analysis

For RNA sequencing (RNA-Seq) library preparation, total RNA was extracted and quality-checked to ensure suitability for library construction. Poly(A)-tailed mRNA was enriched using oligo(dT) magnetic beads and fragmented. First-strand cDNA was synthesized using reverse transcriptase and random primers, followed by second-strand cDNA synthesis. Double-stranded cDNA underwent end repair, adenylation, and adapter ligation. Adapter-ligated cDNA was PCR-amplified, and the library was quality-checked. Single-stranded circular DNA was generated, and DNA nanoballs (DNBs) were produced via rolling circle replication. DNBs were loaded onto high-density DNA nanochips and sequenced using combinatorial probe anchor synthesis (cPAS). Sequencing services were provided by BGI (Wuhan, China). For data analysis, transcriptomic data were analyzed using BGI’s Dr.TOM platform. Differentially expressed genes (DEGs) were identified, and functional enrichment analysis of DEGs was performed using Gene Ontology (GO) and Kyoto Encyclopedia of Genes and Genomes (KEGG) databases. Gene Set Enrichment Analysis (GSEA) was further applied to identify biological processes significantly enriched across sample groups. These analyses provided insights for subsequent experimental validation and mechanistic investigation.

### Untargeted lipidomics

Lipids were extracted from frozen mouse heart tissue (50 mg) by homogenization in 500 μL of chloroform:methanol (2:1, v/v) on ice. After adding water (1:4, v/v), the mixture was vortexed, centrifuged at 3,000 rpm for 15 min, and the organic phase was collected. The solvent was evaporated under nitrogen gas to form a lipid film, which was stored at -80°C for analysis. For lipid profiling by LC-MS/MS, lipids were separated using a Cortes C18 column (2.1 × 100 mm, 2.7 μm) with a gradient of mobile phase A (0.77 g/L ammonium acetate in water/acetonitrile, 4:6, v/v) and mobile phase B (acetonitrile/isopropanol, 1:9, v/v) at 0.25 mL/min. Detection was performed using a Q Exactive™ HF Orbitrap™ mass spectrometer coupled with UHPLC, operating in positive/negative ion modes. Lipid identification and quantification were conducted using LipidSearch software (Thermo Fisher Scientific).

### Untargeted Metabolomics

Metabolites were extracted from frozen mouse heart tissue (50 mg) by homogenization in 500 μL of ice-cold 80% (v/v) HPLC-grade methanol using a handheld electric homogenizer (2–3 cycles, ≤10 s per cycle, on ice). The homogenate was vortexed for 1 min at 4–8°C, incubated overnight at -80°C, and centrifuged (14,000 × *g*, 20 min, 4°C). The supernatant was dried using a vacuum concentrator at room temperature and stored at -80°C until analysis. Metabolite profiling was performed using LC-MS/MS on a TSQ Quantis™ triple quadrupole mass spectrometer coupled to a C18 reversed-phase column. Mobile phases included (A) 10 mM tributylamine/15 mM acetate aqueous solution, and (B) methanol, with a linear gradient of 5–90% B over 25 min. Data were acquired in positive/negative ion switching mode (Q1/Q3 resolution: 0.7 FWHM). Key mass spectrometry parameters were as follows: ion spray voltage, 3.5 kV (positive) and 2.5 kV (negative); capillary temperature, 320°C; sheath gas pressure, 35 arb; auxiliary gas pressure, 10 arb. Metabolites were identified using Tracefinder™ software (Thermo Fisher Scientific).

### Isolation and culture of primary neonatal mouse cardiomyocytes

Primary neonatal mouse cardiomyocytes (NMCMs) were isolated from 1-day-old WT pups (n=30–40). Hearts were excised under sterile conditions after disinfection in 75% ethanol, minced, and digested sequentially in pre-warmed 0.1% trypsin (T6567, Sigma) and 0.08% collagenase II (LS004176, Worthington). Both enzymes were prepared in HBSS (311-512, Wisent) with 10 mM HEPES (H810752, Macklin). Digestion was performed on a shaking table at 37°C. Digested tissue fragments were filtered through a 70-μm cell strainer (Falcon), centrifuged (1,000 rpm, 5 min), and resuspended in DMEM/F12 culture medium (319-085, Wisent) containing 10% fetal bovine serum (FBS; Vivacell), 100 U/mL penicillin and 100 μg/mL streptomycin (450-201, Wisent). To enrich for cardiomyocytes, the cell suspension was subjected to differential adhesion for 90 min on dishes pre-coated with 1% gelatin (ST1339, Beyotime). The non-adherent cardiomyocytes were collected, resuspended in DMEM/F12 medium containing 0.1 mM 5-bromo-2’-deoxyuridine (BrdU; ST1056, Beyotime), and seeded at densities of 5–6×10⁵ cells/well in 6-well plates for molecular analyses or 1–2×10⁵ cells/dish in 35-mm glass-bottom culture dishes (NEST) that were pre-coated with gelatin for morphological studies. All procedures were performed using sterile buffers and reagents, which were filtered through 0.22-μm filters (SLGP033RB, Millipore) prior to use.

### Treatments of primary cardiomyocytes

Following a 24-h serum starvation, NMCMs were treated with 5 mg/mL rMSA in serum-free medium for 24 h, and control cells received an equal volume of ddH₂O. Cells were then harvested for subsequent analyses, such as qRT-PCR, and fatty acid oxidation assays.

To investigate the signaling pathways, serum-starved NMCMs were treated with either varying concentrations of rMSA for 6 h or with 5 mg/mL rMSA for different durations. In some experiments, specific pathways were inhibited: AMPK activity was suppressed using 5 μM Dorsomorphin dihydrochloride (Compound C; S7306, Selleck), and PKA activity was suppressed using 5 μM H-89 dihydrochloride (S1582, Selleck), with both inhibitors applied for 24 h prior to rMSA treatment.

Transfection was performed using the Polyplus jetPRIME kit (101000046, Polyplus). Mouse-specific *Lkb1* siRNA (sense: 5′-GCGCCAAGCUCAUCGGCAATT-3′; antisense: 5′-UUGCCGAUGAGCUUGGCGCTT-3′) and a negative control siRNA (sense: 5′-UUCUCCGAACGUGUCACGUTT-3′; antisense: 5′-ACGUGACACGUUCGGAGAATT-3′) were obtained from Genepharma. After 48 h of transfection in DMEM/F12 complete medium, the medium was replaced with serum-free medium, followed by rMSA treatment.

To compare the efficacy of albumin from different sources, plasma-derived mouse serum albumin (pMSA; CSB-NP00080, Cusabio) was used at the same concentration as rMSA in some parallel experiments.

To establish an *in vitro* model of cardiomyocyte hypertrophy, NMCMs were serum-starved for 24 h and then stimulated with 200 µM phenylephrine (PE; S2569, Selleck) in serum-free medium for 48 h. To evaluate the interventional effect of rMSA on induced hypertrophy, cells were first treated with 200 µM PE for 24 h, followed by the addition of 5 mg/mL rMSA, and control cells received an equal volume of ddH₂O. Depending on the experimental aims, this PE-induced hypertrophy paradigm was combined with pharmacological inhibitors, siRNA transfection, or pMSA intervention. Cells were collected or assayed according to the defined experimental endpoints.

### Immunofluorescence (IF) staining

#### Cardiac tissue sections

Paraformaldehyde-fixed cardiac tissues were dehydrated in a sucrose gradient, embedded in OCT compound, and sectioned at 5-μm thickness using a cryostat. Sections were equilibrated to room temperature, permeabilized with 0.1% Triton X-100 (P0096, Beyotime) for 10 min, and washed three times with PBS. Non-specific binding sites were blocked with 10% goat serum (ZLI-9056, ZSGB-Bio) for 1 h at room temperature. Sections were then incubated overnight at 4°C with the anti-albumin primary antibody (1:200, 16475-1-AP, Proteintech). After washing with PBS, sections were incubated with the secondary antibody and WGA for 1 h at room temperature in the dark, followed by nuclear counterstaining with DAPI (1:1000, C0060, Solarbio) for 3 min. Stained sections were visualized using a confocal microscope.

#### Primary cardiomyocytes

Cells cultured on 35-mm glass-bottom culture dishes (Nest) were fixed in acetone:methanol (1:1, v/v) at 4°C for 10 min, permeabilized with 0.1% Triton X-100, and blocked with 10% goat serum. Primary antibodies against α-actinin (1:500, A7811, Sigma-Aldrich) and/or albumin (1:200, 16475-1-AP, Proteintech) were applied overnight at 4°C. After secondary antibody incubation and DAPI staining, IF imaging was performed using a confocal microscope.

### Metabolic experiments on isolated mitochondria

#### Mitochondria isolation

Mitochondria were isolated from adult mouse left ventricular tissues (approximately 50 mg) using a tissue mitochondria isolation kit (C3606, Beyotime), with all procedures performed on ice. Briefly, tissues were rinsed with ice-cold PBS, minced, and digested with ice-cold trypsin for 20 min. After a brief centrifugation (600 × *g*, 10–20 s), the pellet was resuspended in Mitochondria Isolation Reagent A and homogenized with 20– 30 strokes. Sequential centrifugations were performed at 1,000 × *g* for 5 min and then at 3,500 × *g* for 10 min to pellet the mitochondria. All centrifugation steps were carried out at 4°C.

#### Mitochondrial respiratory function experiment of isolated mitochondria from cardiac tissues

Mitochondrial respiratory function was assessed using the Seahorse XFe96 Analyzer (Agilent). A 1× mitochondrial assay solution (MAS, pH 7.2 at 37°C) containing 70 mM sucrose (S818049, Macklin), 220 mM mannitol (M6266, Macklin), 10 mM KH₂PO₄ (P815660, Macklin), 5 mM MgCl₂ (M872005, Macklin), 2 mM HEPES (H810752, Macklin), 1 mM EGTA (E6257, Macklin), and 0.2% (w/v) fatty acid-free BSA (ST025, Beyotime) was prepared and stored at -20°C. Substrate stocks which were 0.5 M succinate (S2378, Sigma), 0.5 M malate (M6413, Sigma), 0.5 M glutamate (G8415, Sigma), 0.5 M pyruvate (S2378, Sigma), and 40 mM ADP (S2378, Sigma) in 1× MAS were pH-adjusted to 7.2, with 0.5 M pyruvate (107360, Sigma) freshly prepared. Respiratory assay reagents included 1 M ascorbate (A92902, Sigma) and 10 mM TMPD (T7394, Sigma) dissolved in 1× MAS at pH 7.2, and DMSO-dissolved stocks which were 5 mM oligomycin (ab141829, Abcam), 2.5 mM FCCP (C2920, Sigma), 2.5 mM antimycin A (ab141904, Abcam), and 2.5 mM rotenone (R8875, Sigma).

For the coupling assay, isolated cardiac mitochondria were resuspended in ice-cold 1× MAS containing 10 mM glutamate and 5 mM malate. Isolated cardiac mitochondria were adjusted to 5 μg/well using BCA assay, seeded onto the Seahorse XFe96 plate (25 μL/well), and centrifuged (2,000 × *g*, 20 min, 4°C), followed by addition of 155 μL pre-warmed (37°C) assay solution. One day prior to testing, the probe plate was hydrated with Seahorse XF Calibrant Solution (100840-000, Agilent) in a CO₂-free incubator at 37°C. A pre-hydrated sensor cartridge was loaded with compounds to achieve the following final concentrations: port A, 4 mM ADP; port B, 2.5 μg/mL oligomycin; port C, 4 μM FCCP; port D, 4 μM antimycin A. Oxygen consumption rates (OCR) were measured after system calibration, with State II (basal), State III (ADP-stimulated), and State IIIu (FCCP-uncoupled) respiration calculated by subtracting non-mitochondrial respiration. Twelve technical replicates per sample were analyzed.

For the electron flow assay, mitochondrial electron transport chain activity was analyzed. 1× MAS containing 10 mM pyruvate, 2 mM malate, and 4 μM FCCP was prepared for mitochondria resuspension. Isolated cardiac mitochondria were adjusted to 3 μg/well using BCA assay, seeded onto the Seahorse XFe96 plate (25 μL/well), and centrifuged (2,000 × *g*, 20 min, 4°C), followed by addition of 155 μL pre-warmed (37°C) assay solution. During centrifugation, a pre-hydrated sensor cartridge was loaded with compounds to achieve the following final concentrations: port A, 2 μM rotenone; port B, 10 mM succinate; port C, 4 μM antimycin A; port D, 10 mM ascorbate / 100 μM TMPD. OCR were measured post-calibration. Complex I-driven respiration was assessed under uncoupled conditions (pyruvate/malate/FCCP), followed by sequential inhibition with rotenone (Complex I blockade), succinate-induced Complex II activity, antimycin A (Complex III inhibition), and ascorbate/TMPD-mediated Complex IV respiration. Twelve technical replicates per sample were analyzed, with OCR values reflecting specific electron transport chain contributions.

#### Mitochondrial stress assay in primary cardiomyocytes

Primary neonatal mouse cardiomyocytes were isolated and seeded onto the Seahorse XFe96 plate at 2.5×10⁴ cells/well. After 3 days of culture, cells were serum-starved for 24 h and then treated with 200 µM PE for 24 h. Subsequently, cells were co-treated with or without rMSA for another 24 h, generating four groups: NC, rMSA, PE, and PE + rMSA. On the day before the assay, the probe plate was hydrated in a CO₂-free incubator at 37°C. The mitochondrial stress assay medium was freshly prepared using Seahorse XF DMEM Medium (103575-100, Agilent) and supplemented with 10 mM glucose (G7528, Sigma), 2 mM L-glutamine (G8540, Sigma), and 1 mM sodium pyruvate (S8636, Sigma). On the assay day, cells were washed three times with mitochondrial stress assay medium and equilibrated in 175 μL/well of fresh medium for 1 h at 37°C (CO₂-free). A pre-hydrated sensor cartridge was loaded with compounds to achieve the following final concentrations: port A, 2 μM oligomycin; port B, 1 μM FCCP; port C, 1 μM antimycin A and 1 μM rotenone. After the probe plate calibration, the cell culture plate was transferred to the Seahorse analyzer for real-time OCR measurement. Key parameters including basal respiration, ATP production, maximal respiration, and spare respiratory capacity were calculated from OCR data after subtraction of non-mitochondrial respiration.

#### Palmitate-induced oxidative stress assay in primary cardiomyocytes

NMCMs were isolated and seeded onto the Seahorse XFe96 culture plate at 2.5×10⁴ cells/well. After 3 days of adherent culture, cells were serum-starved in DMEM/F12 medium for 24 h. Experimental groups were treated with 5 mg/mL rMSA for 16 h, while controls received an equal volume of ddH₂O, resulting in four groups: Vehicle-Ctrl, Vehicle-rMSA, Palmitate-Ctrl, and Palmitate-rMSA. 4 h prior to the assay, the medium was replaced with a substrate-limited medium (low-glucose DMEM containing 0.5 mM carnitine (HY-B2246, MCE)). For the assay, this medium was replaced with freshly prepared FAO assay medium (Seahorse XF DMEM Medium containing 2.5 mM glucose and 0.5 mM carnitine). Cells were equilibrated in FAO medium (150 μL/well) at 37°C in a CO₂-free incubator for 1 h. A sensor cartridge was hydrated overnight under the same conditions. Simultaneously, a pre-hydrated sensor cartridge was loaded with compounds to achieve the following final concentrations: port A, 2 μM oligomycin; port B, 3 μM FCCP; port C, 1 μM antimycin A and 1 μM rotenone. At Time 0, 3.5 μL of either 7.5 mM palmitate conjugated to BSA (Palm-BSA; for palmitate groups) or 1.5 mM BSA (for vehicle groups) was added to each well, ensuring a final BSA concentration of 150 μM. A 7.5 mM palmitate-BSA (Palm-BSA) stock solution was prepared as follows: First, a 250 mM sodium palmitate (S85020, Acmec) solution was prepared in 50% ethanol. Then, 30 μL of this 250 mM sodium palmitate-ethanol solution was rapidly added to 1 mL of pre-warmed (37°C) 1.5 mM BSA stock solution to obtain the final 7.5 mM Palm-BSA conjugate. After the probe plate calibration, OCR was measured, and basal and maximal respiration were calculated from OCR data after subtraction of non-mitochondrial respiration.

#### Mitochondrial membrane potential assay

The mitochondrial membrane potential was assessed using an Enhanced JC-1 Assay Kit (C2003S, Beyotime). Cells cultured on glass-bottom dishes were washed with PBS and incubated with 1 mL JC-1 working solution at 37°C for 20 min. After two washes with JC-1 staining buffer, fresh culture medium was added for imaging. Fluorescence was visualized using a confocal laser scanning microscope. JC-1 monomers (excitation/emission: 490/530 nm) and aggregates (excitation/emission: 525/590 nm) were imaged. The ratio of aggregate (red) to monomer (green) fluorescence intensity, quantified using ImageJ software, served as a relative indicator of mitochondrial membrane potential.

### Statistical analysis

All statistical metrics and significance annotations were provided in both the figures and their respective legends. The cell experimental data in this study were obtained through at least three independent biological replicates to ensure result reliability and reproducibility. Data were presented as the mean ± SEM unless otherwise stated, and statistical analyses were performed using GraphPad Prism software (version 10). For comparisons between two groups, an unpaired two-tailed Student’s *t*-test was applied. Multi-group comparisons involving multiple factors were analyzed using two-way ANOVA followed by Tukey’s multiple comparisons test for inter-group difference assessment. For comparisons among three or more groups, one-way ANOVA followed by Tukey’s multiple comparisons test was used. All statistical tests were preceded by validation of normality and homogeneity of variance assumptions, with the significance threshold set at *p*<0.05. Differences were annotated using standardized notation: ns (not significant), \**p*<0.05, \*\**p*<0.01, \*\*\**p*<0.001, and \*\*\*\**p*<0.0001. Significance markers in figures were strictly derived from statistical outcomes, with error bars consistently representing SEM ranges throughout the study.

## Acknowledgments

We thank Shenzhen Protgen, Ltd. for generously supplying rMSA. We thank the Center of Biomedical Analysis of Tsinghua University for Seahorse tests; the State Key Laboratory of Membrane Biology for confocal microscopy; the Cell Biology Facility of the Center of Biomedical Analysis of Tsinghua University for TEM; the Metabolomics Facility Center of Metabolomics and Lipidomics in National Protein Science Technology Center of Tsinghua University for LC-MS/MS. We thank the Luo laboratory members for their technical assistance and valuable insights on this study.

## Author contributions

Conceptualization: Q.A., B.M.; Methodology: Q.A., B.M.; Investigation: Q.A.; Software: Q.A.; Writing & Editing & Reviewing: Q.A., B.M., Y.F., Y.L.; Supervision: Y.L. All authors have read and approved the article.

