## Supplementary Figures and tables for "Albumin alleviates pressure overload-induced heart failure by improving mitochondrial homeostasis via PKA-LKB1-AMPK axis"

**Figure S1**

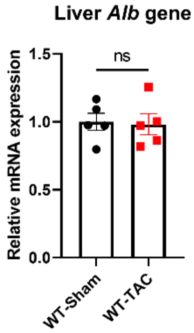


**Figure S1.** ***Alb*** **mRNA level in liver tissues**.

mRNA expression of *Alb* in liver tissues from WT mice after sham or TAC 6 weeks, normalized to *Actb* mRNA (n = 5). Mean ± SEM, unpaired two-tailed Student's *t*-test. ns, not significant.

#### Figure S2

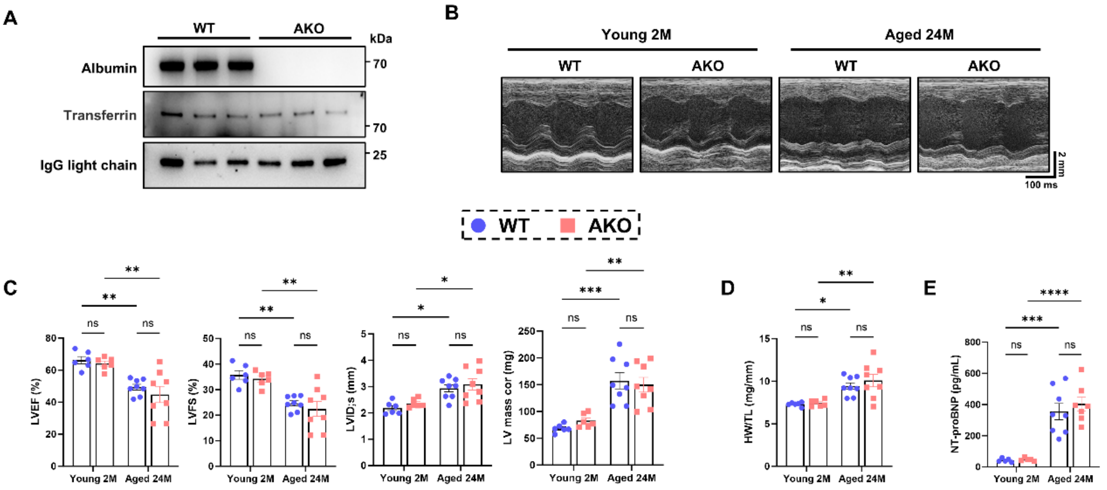


**Figure S2. Genetic albumin deficiency does not exacerbate cardiac dysfunction or remodeling in young or aged mice.**

(A) Representative immunoblots of serum albumin levels from WT and AKO mice, with serum transferrin and IgG light chain as endogenous controls (n = 3).

(B and C) Representative M-mode echocardiographs (B) and quantification (C) of LVEF, LVFS, LVID;s and LV mass cor in young (2 months, 2M) or aged (24 months, 24M) WT and AKO mice (n = 6–8). Scale bars, vertical 2 mm and horizontal 100 ms.

(D) Quantification of HW/TL in young or aged WT and AKO mice (n = 6–8).

(E) Serum NT-proBNP levels of young or aged WT and AKO mice (n = 5–8).

Mean ± SEM, two-way ANOVA followed by Tukey's multiple comparisons test. ns, not significant; **p*<0.05, ***p*<0.01, ****p*<0.001, *****p*<0.0001.

**Figure S3**

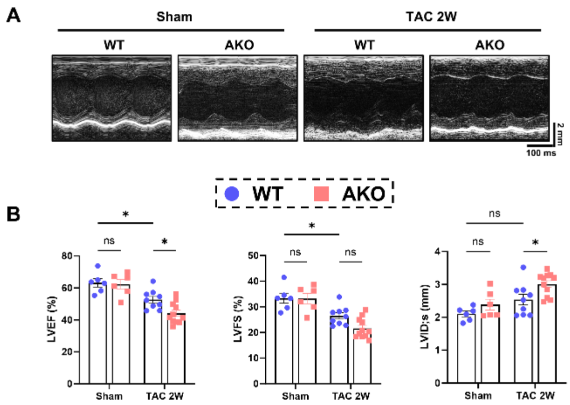


**Figure S3. Echocardiography in WT or AKO mice after sham or TAC 2 weeks.** Representative M-mode echocardiographs (A) and quantification (B) of LVEF, LVFS and LVID;s in WT and AKO mice after sham or TAC 2 weeks (n = 6–11). Scale bars, vertical 2 mm and horizontal 100 ms. Mean ± SEM, two-way ANOVA followed by Tukey's multiple comparisons test. ns, not significant; *$p<0.05$.

### Figure S4

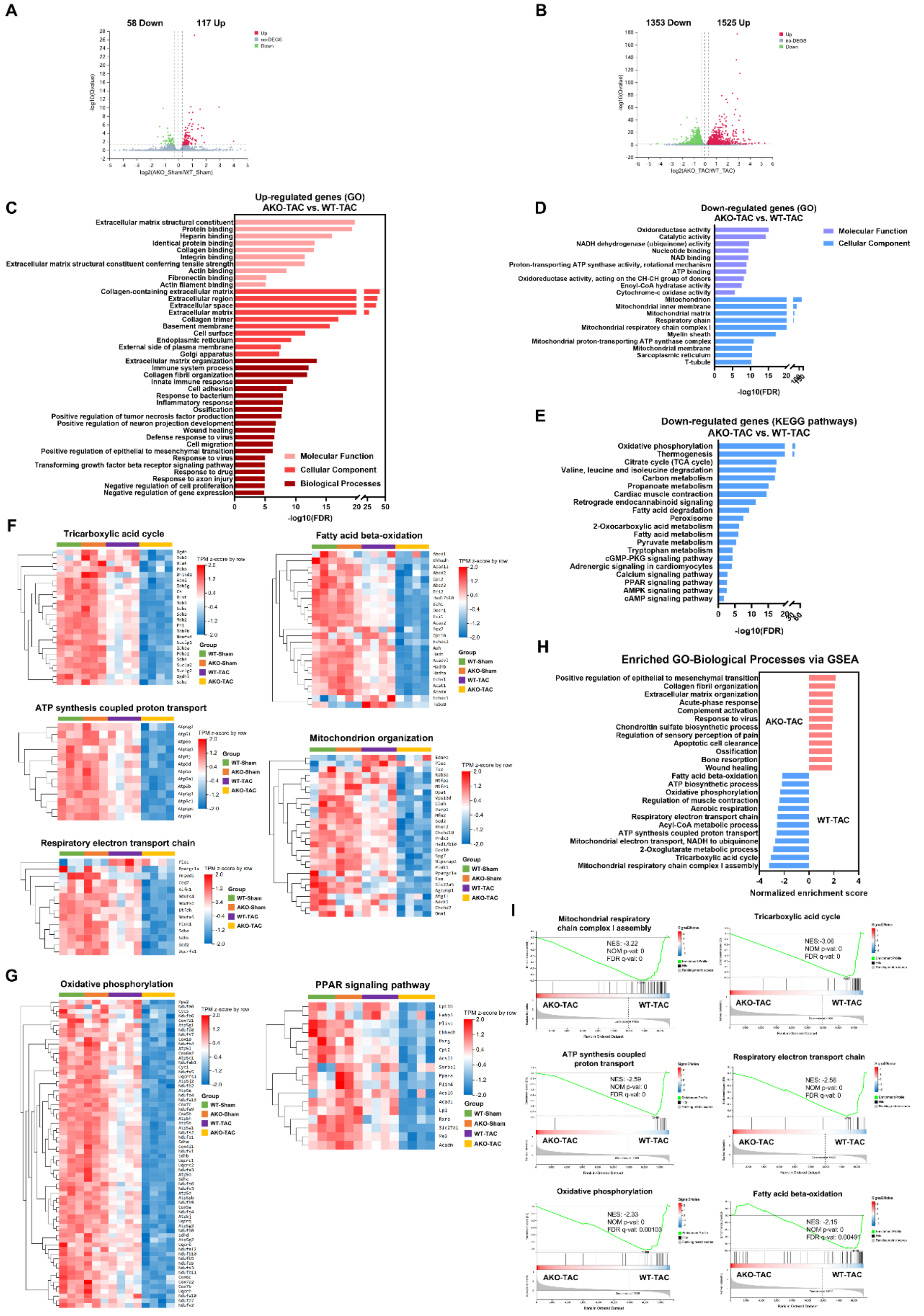


**Figure S4. RNA sequencing in cardiac tissues from WT or AKO mice after sham or TAC 6 weeks.**

(A) Volcano plot depicting DEGs of cardiac tissues from AKO-Sham and WT-Sham groups (n = 3).

(B) Volcano plot depicting DEGs of cardiac tissues from WT and AKO mice after TAC 6 weeks (n = 4).

(C and D) GO enrichment analysis of molecular function, cellular component and biological processes for genes significantly up-regulated in AKO-TAC group (vs. WT-TAC) (C, n = 4) or molecular function and cellular component for genes significantly down-regulated in AKO-TAC group (vs. WT-TAC) (D, n = 4).

(E) KEGG pathway enrichment analysis of genes significantly down-regulated in the AKO-TAC group (vs. WT-TAC) (n = 4).

(F and G) Heatmaps displaying transcriptomic profiles of representative GO biological processes (F) and KEGG pathways (G) across WT-Sham, AKO-Sham, WT-TAC, and AKO-TAC groups (n = 3–4). Genes down-regulated in AKO-TAC (vs. WT-TAC) were visualized.

(H) GSEA of GO biological processes in cardiac tissues from AKO-TAC and WT-TAC groups (n = 4). Significantly enriched terms (|NES| > 1, NOM p-val < 0.05, FDR q-val < 0.25) were shown.

(I) GSEA enrichment plots of representative biological processes significantly enriched in WT-TAC group (n = 4). Key statistics (NES, NOM p-val, FDR q-val) were annotated.

(A and B) Red and green data points represented significantly up- and down-regulated genes, respectively. Transcripts (|fold change| > 1.2, Q value (adjusted p-value) < 0.05) were shown.

(C–E) Representative enriched terms (FDR < 0.05) were ranked by significance (-$\log_{10}$FDR). FDR, false discovery rate (*Q* value).

#### Figure S5

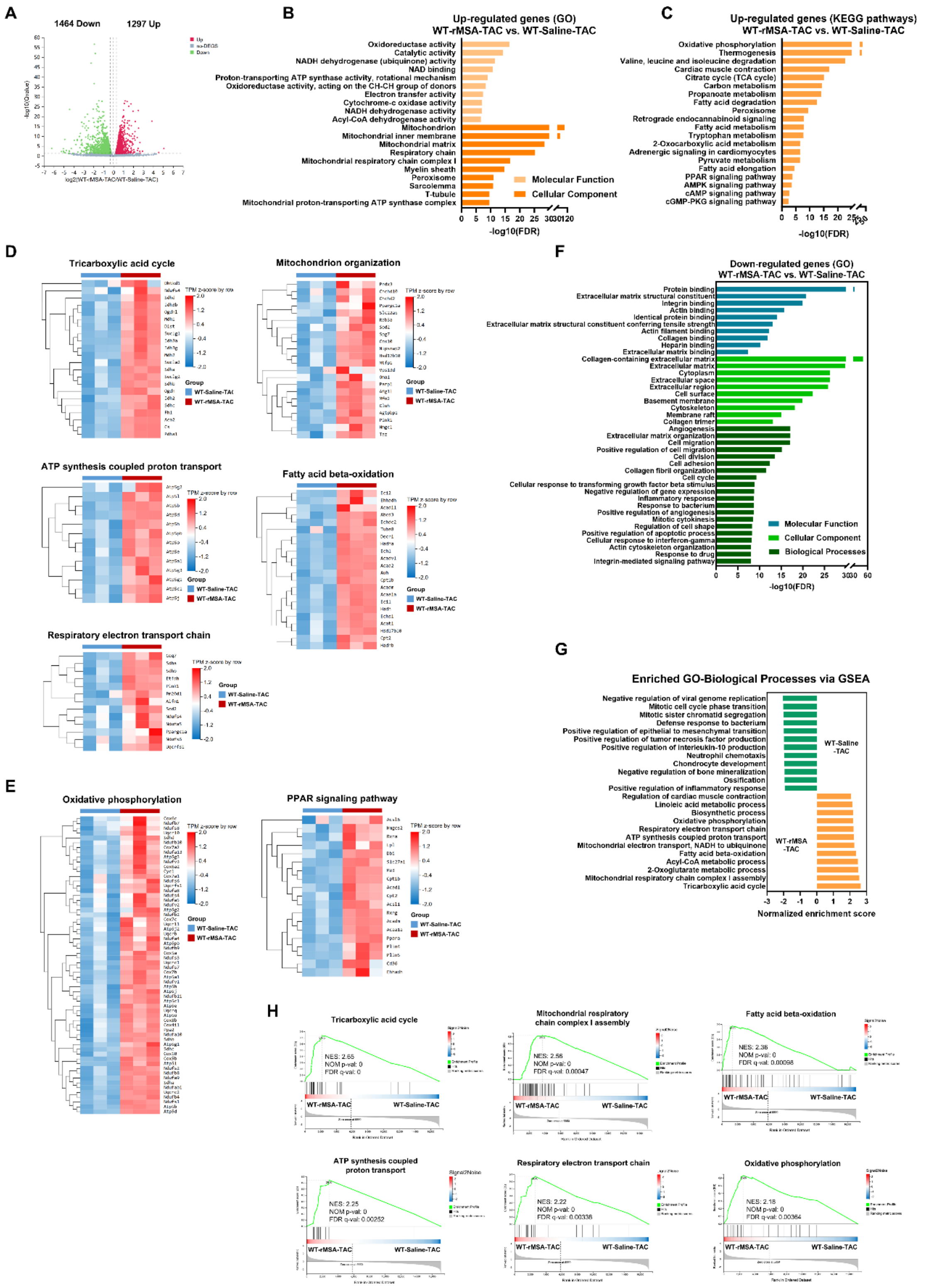


**Figure S5. RNA sequencing in cardiac tissues from WT-Saline or WT-rMSA mice after sham or TAC 8 weeks.**

(A) Volcano plot depicting DEGs of cardiac tissues from WT-Saline-TAC and WT-rMSA-TAC groups (n = 3). Red and green data points represented significantly up- and down-regulated genes, respectively. Transcripts (|fold change| > 1.2, Q value (adjusted p-value) < 0.05) were shown.

(B) GO enrichment analysis of molecular function and cellular component for genes significantly up-regulated in WT-rMSA-TAC group (vs. WT-Saline-TAC) (n = 3).

(C) KEGG pathway enrichment analysis of genes significantly up-regulated in WT-rMSA-TAC group (vs. WT-Saline-TAC) (n = 3).

(D and E) Heatmaps displaying transcriptomic profiles of representative GO biological processes (D) and KEGG pathways (E) across WT-rMSA-TAC and WT-Saline-TAC groups (n = 3). Genes up-regulated in WT-rMSA-TAC group (vs. WT-Saline-TAC) were visualized.

(F) GO enrichment analysis of molecular function, cellular component and biological processes for genes significantly down-regulated in WT-rMSA-TAC group (vs. WT-Saline-TAC) (n = 3).

(G) GSEA of GO biological processes in cardiac tissues from WT-rMSA-TAC and WT-Saline-TAC groups (n = 3). Significantly enriched terms (|NES| > 1, NOM p-val < 0.05, FDR q-val < 0.25) were shown.

(H) GSEA enrichment plots of representative biological processes significantly enriched in WT-rMSA-TAC group (n = 3). Key statistics (NES, NOM p-val, FDR q-val) were annotated.

(B, C and F) Representative enriched terms (FDR < 0.05) were ranked by significance ($-\log_{10}$FDR). FDR, false discovery rate (*Q* value).

**Figure S6**

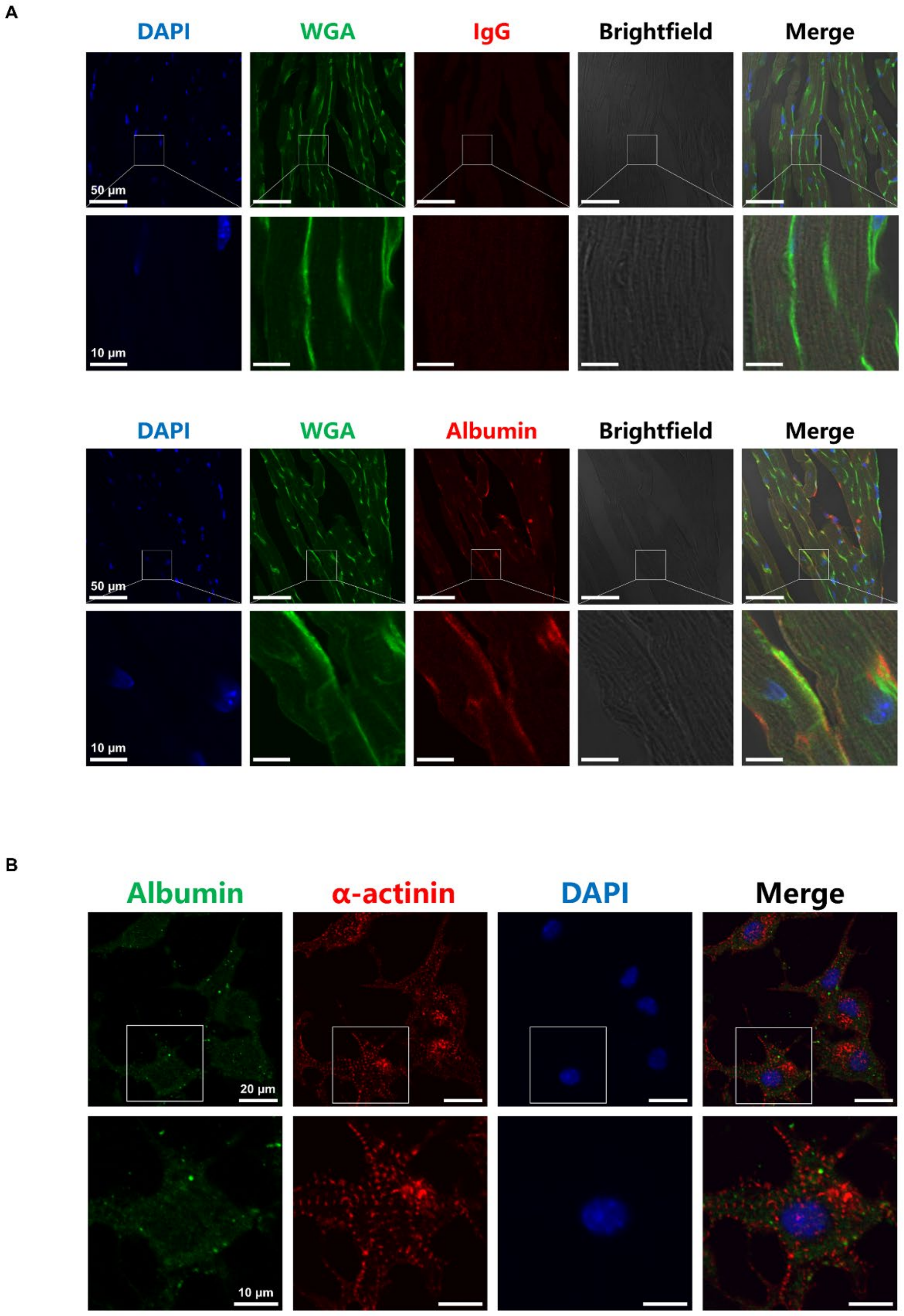


**Figure S6. Albumin can contact cardiomyocytes and be endocytosed into the cells.** Representative immunofluorescence staining of albumin in cardiac tissues from WT mice (A), and primary cardiomyocytes (B). In (A), WGA was used to outline cardiomyocyte borders, while "IgG" indicates control staining with the secondary antibody alone. In (B), α-Actinin was employed to identify the isolated primary cardiomyocytes. Scale bars are shown in the panels.

### Figure S7

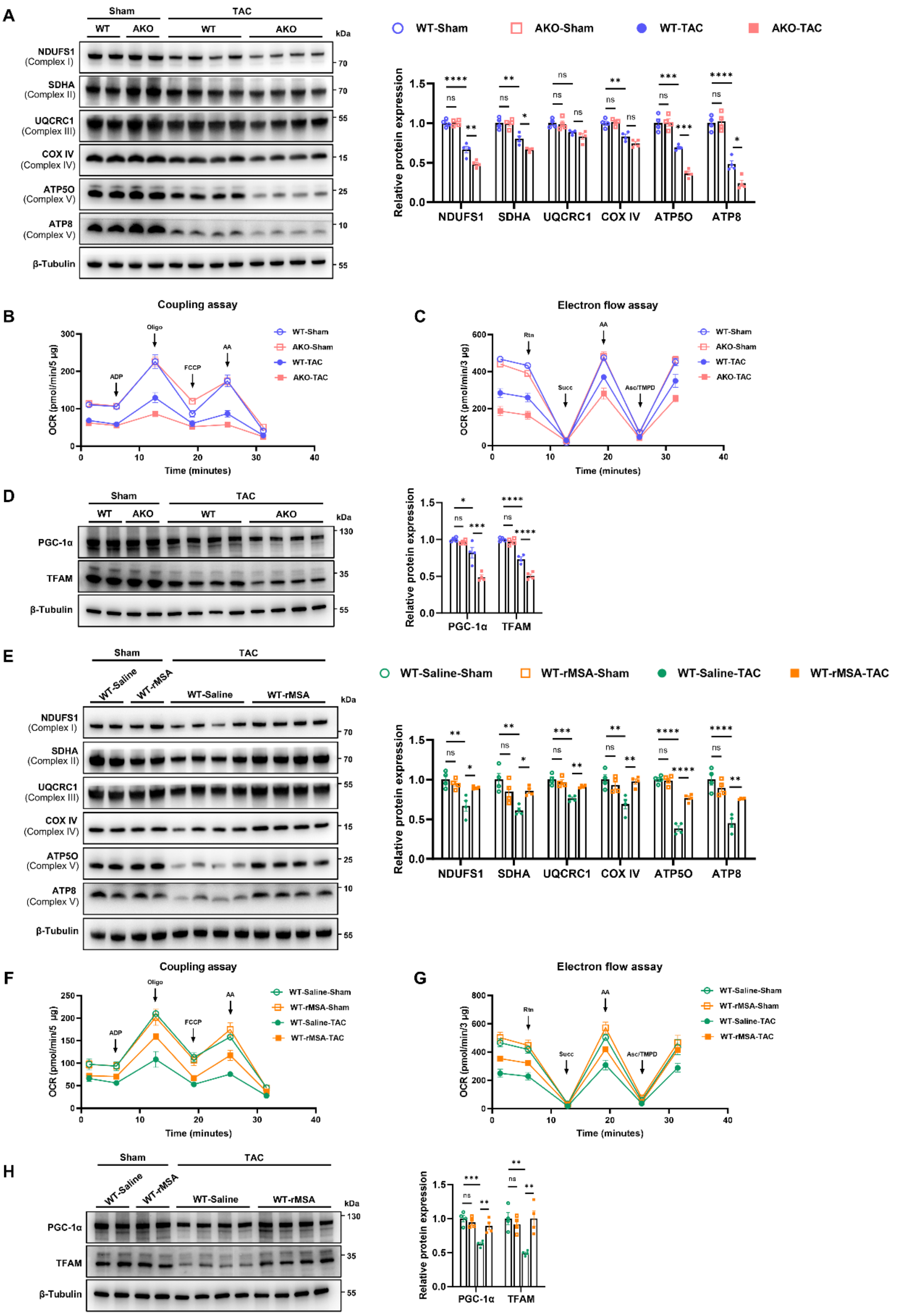


**Figure S7. Albumin enhances cardiac mitochondrial respiratory function and preserves MQC in pressure overload-induced heart failure.**

(A–D) Samples from WT and AKO mice after sham or TAC 6 weeks.

(A) Representative immunoblots and quantitation of ETC proteins in cardiac tissues (n = 4).

(B and C) Measurement of OCR in the coupling assay (B, n = 4–8), and in the electron flow assay (C, n = 5–6) in isolated cardiac mitochondria.

(D) Representative immunoblots and quantitation of PGC-1α and TFAM proteins in cardiac tissues (n = 4).

(E–H) Samples from WT mice treated with rMSA or saline after sham or TAC 8 weeks.

(E) Representative immunoblots and quantitation of ETC proteins in cardiac tissues (n = 4).

(F and G) Measurement of OCR in the coupling assay (F, n = 4–5), and in the electron flow assay of isolated cardiac mitochondria (G, n = 4–7).

(H) Representative immunoblots and quantitation of PGC-1α and TFAM proteins in cardiac tissues (n = 4).

Key subunits from mitochondrial ETC complexes including NDUFS1 (Complex I), SDHA (Complex II), UQCRC1 (Complex III), COX IV (Complex IV), and ATP5O and ATP8 (Complex V) were detected in (A) and (E), and β-Tubulin served as the loading control.

(B and F) Complex I-driven respiration was initiated with 10 mM glutamate and 5 mM malate. OCR was measured under sequential conditions: basal OCR (State II), followed by injections of ADP (4 mM, State III), oligomycin (2.5 μg/mL), FCCP (4 μM, State IIIu), and antimycin A (4 μM, Non-mitochondrial OCR). Non-mitochondrial (Non-mito) OCR was subtracted from State II, State III and State IIIu OCR values. Mitochondria (5 μg/well, 12 technical replicates/mouse).

(C and G) OCR was measured under sequential conditions: pyruvate/malate/FCCP (10 mM/2 mM/4 μM) to uncouple mitochondria and assess maximal Complex I-driven respiration; 2 μM rotenone to inhibit Complex I; 10 mM succinate to activate Complex II-driven respiration; 4 μM antimycin A to inhibit Complex III; and ascorbate/TMPD (10 mM/100 μM) to measure Complex IV-driven respiration. Mitochondria (3 μg/well, 12 technical replicates/mouse).

Mean ± SEM, two-way ANOVA followed by Tukey's multiple comparisons test. ns, not significant; *$p$<0.05, **$p$<0.01, ***$p$<0.001, ****$p$<0.0001.

**Figure S8**

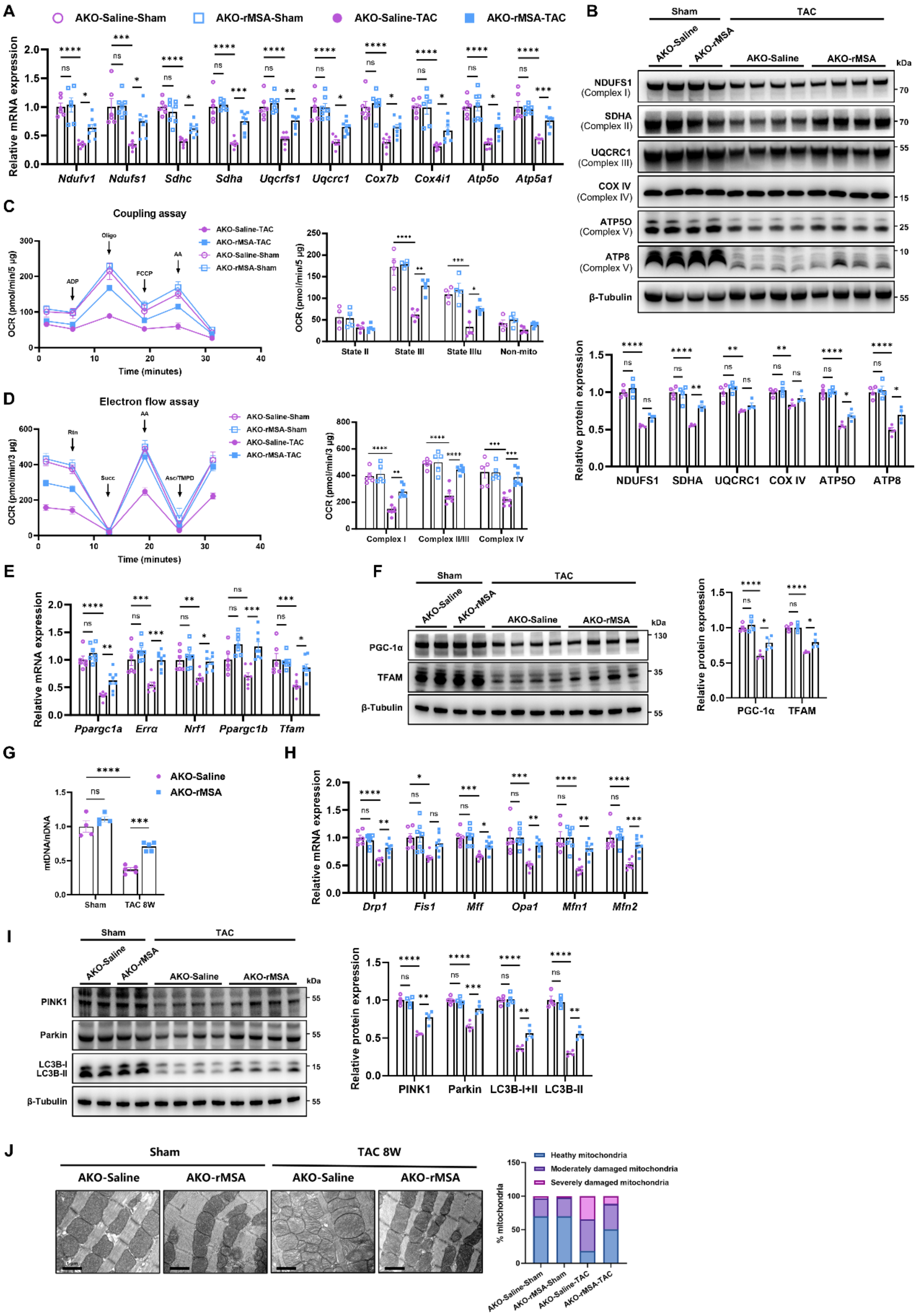


**Figure S8. Albumin attenuates the progression of cardiac mitochondrial respiratory dysfunction and preserves MQC in pressure overload-induced heart failure AKO mice.**

Analyses were performed in AKO mice treated with rMSA or saline after sham or TAC 8 weeks.

(A) mRNA expression of cardiac ETC genes (n = 6–8).

(B) Representative immunoblots and quantitation of ETC proteins in cardiac tissues (n = 4).

(C) Measurement and quantification of the coupling assay in isolated cardiac mitochondria (n = 4–5).

(D) Measurement and quantification of the electron flow assay in isolated cardiac mitochondria (n = 5–7).

(E) Cardiac mRNA expression of mitochondrial biogenesis-associated genes (n = 6–8).

(F) Representative immunoblots and quantitation of PGC-1α and TFAM proteins in cardiac tissues (n = 4).

(G) mtDNA/nDNA ratio in cardiac tissues (n = 4–5).

(H) Cardiac mRNA expression of mitochondrial fission- and fusion-associated genes (n = 6–8).

(I) Representative immunoblots and quantitation of mitophagy-associated proteins in cardiac tissues (n = 4).

(J) Representative TEM images (scale bar, 1 μm) and quantitative classification of mitochondrial morphology in cardiac tissues (n = 4). Mitochondria from random fields (>200 mitochondria per mouse) were classified as healthy, moderately damaged, or severely damaged.

Mean ± SEM, two-way ANOVA followed by Tukey's multiple comparisons test. ns, not significant; *$p<0.05$, **$p<0.01$, ***$p<0.001$, ****$p<0.0001$. mtDNA, mitochondrial DNA; nDNA, nuclear DNA.

**Figure S9**

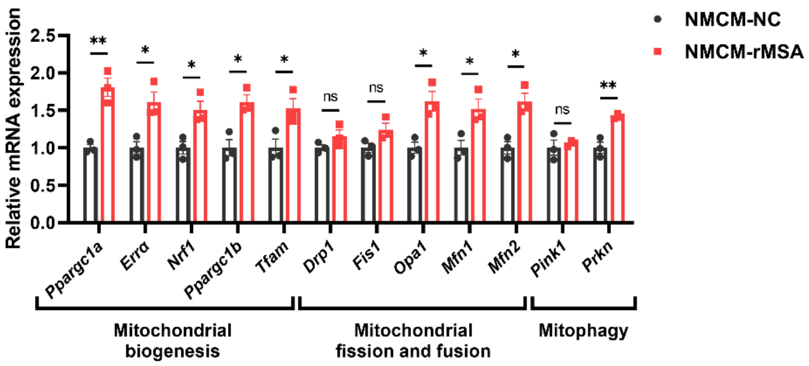


**Figure S9. rMSA treatment affects the mRNA levels of mitochondrial quality control-associated genes.**

qRT-PCR was performed on NMCMs treated with 5 mg/mL rMSA (NMCM-rMSA) or an equal volume of $ddH_2O$ (NMCM-NC). mRNA expression of mitochondrial biogenesis-associated genes (*Ppargc1a*, *Errα*, *Nrf1*, *Ppargc1b* and *Tfam*), mitochondrial fission-associated genes (*Drp1* and *Fis1*) and fusion-associated genes (*Opa1*, *Mfn1* and *Mfn2*), and mitophagy-associated genes (*Pink1* and *Prkn*) in NMCMs treated with or without rMSA, normalized to *Actb* mRNA (n = 3). Mean ± SEM, unpaired two-tailed Student's *t*-test. ns, not significant; *$p$<0.05, **$p$<0.01.

**Figure S10**

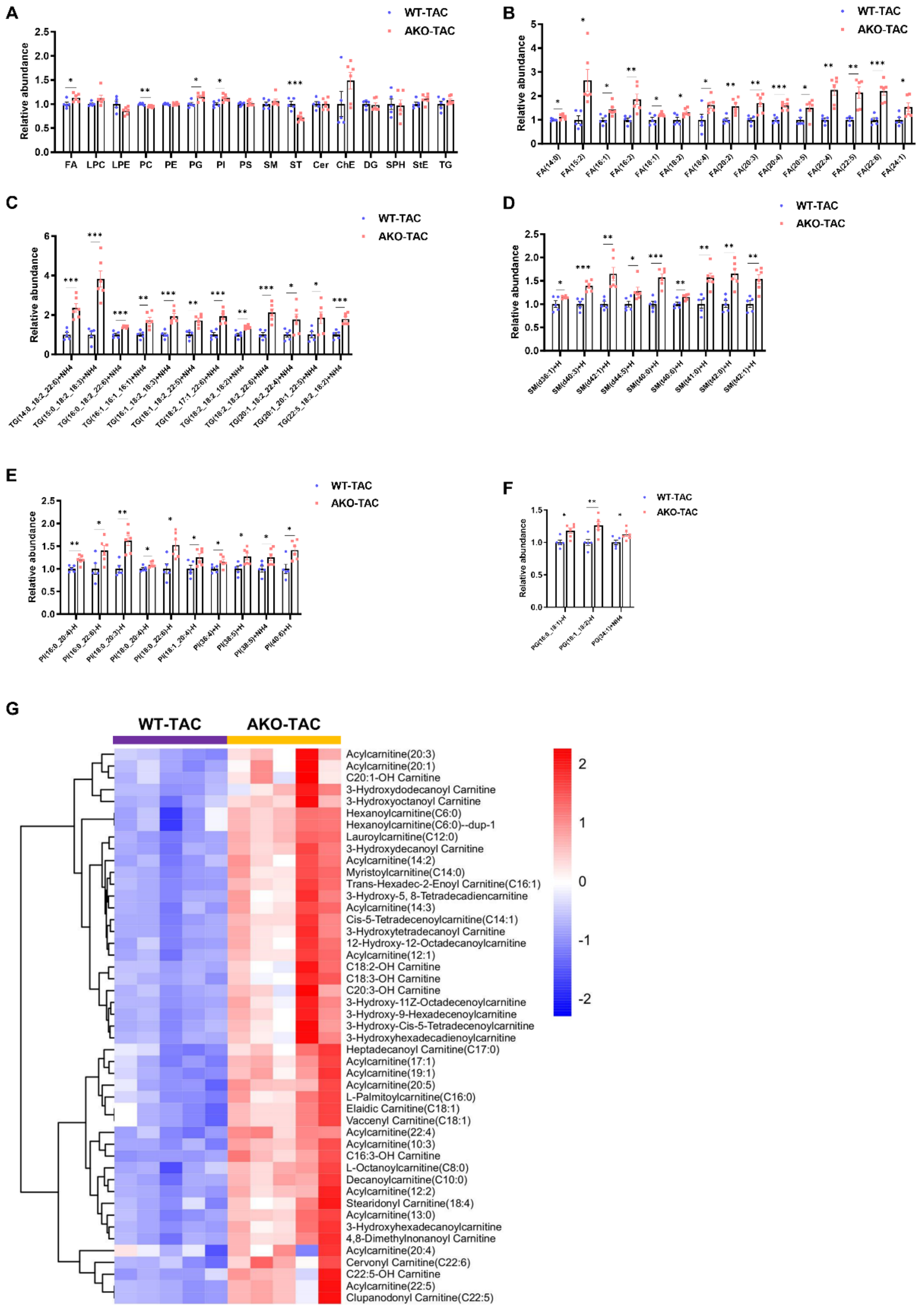


**Figure S10. Cardiac tissue lipidomic and metabolomic profiling in AKO-TAC and WT-TAC mice.**

(A) Comparative analysis of abundances of major lipid classes in cardiac tissues between AKO-TAC and WT-TAC groups after TAC 6 weeks (n = 5–6). Analyzed lipid classes including free fatty acids (FA), lysophosphatidylcholine (LPC), lysophosphatidylethanolamine (LPE), phosphatidylcholine (PC), phosphatidyl ethanolamine (PE), phosphatidylglycerol (PG), phosphatidylinositol (PI), phosphatidylserine (PS), sphingomyelin (SM), sterol lipids (ST), ceramide (Cer), cholesterol ester (ChE), diacylglycerol (DG), sphingosine (SPH), sterol ester (StE), and triacylglycerol (TG).

(B–F) Comparative analysis of significantly altered lipid species including FA (B), TG (C), SM (D), PI (E), PG (F) in AKO-TAC versus WT-TAC groups (n = 5–6).

(G) Heatmap of the comparative abundance of acylcarnitines in cardiac tissues of AKO-TAC vs. WT-TAC groups (n = 6).

Mean ± SEM, unpaired two-tailed Student's *t*-test. ns, not significant; *$p<0.05$, **$p<0.01$, ***$p<0.001$, ****$p<0.0001$.

**Figure S11**

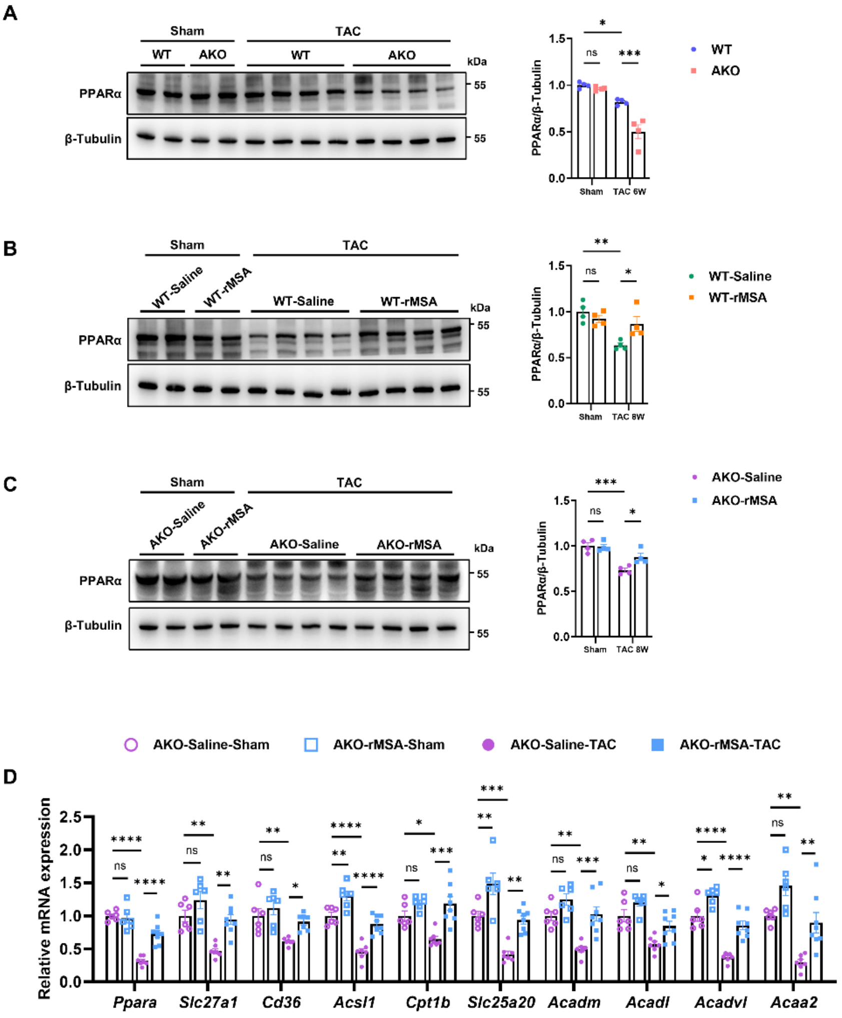


**Figure S11. Albumin enhances the protein levels of PPARα, and rMSA restores FA metabolism gene expression in TAC-induced AKO mice**.

(A–C) Representative immunoblots and quantification of PPARα protein in cardiac tissues from WT and AKO mice after sham or TAC 6 weeks (A, n = 4), from WT mice treated with rMSA or saline after sham or TAC 8 weeks (B, n = 4), and from AKO mice treated with rMSA or saline after sham or TAC 8 weeks (C, n = 4). β-Tubulin served as the loading control.

(D) mRNA expression of cardiac fatty acid metabolism-associated genes in cardiac tissues from AKO mice treated with rMSA or saline after sham or TAC 8 weeks (n = 6–8).

Mean ± SEM, two-way ANOVA followed by Tukey's multiple comparisons test. ns, not significant; *$p<0.05$, **$p<0.01$, ***$p<0.001$, ****$p<0.0001$.

**Figure S12**

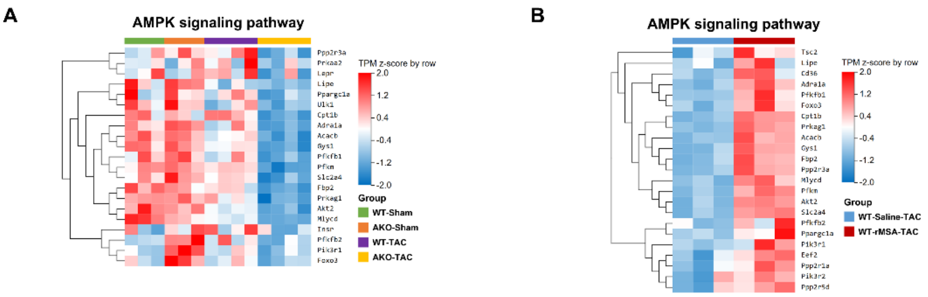


**Figure S12. Heatmaps of genes involved in AMPK signaling pathway.**

Heatmaps of AMPK signaling pathway transcriptomic profiles from cardiac tissues of WT-Sham, AKO-Sham, WT-TAC, and AKO-TAC groups, visualizing genes down-regulated in AKO-TAC (vs. WT-TAC) (A, n = 3–4), and from cardiac tissues of WT-Saline-TAC and WT-rMSA-TAC groups, visualizing genes up-regulated in WT-rMSA-TAC group (vs. WT-Saline-TAC) (B, n = 3).

**Figure S13**

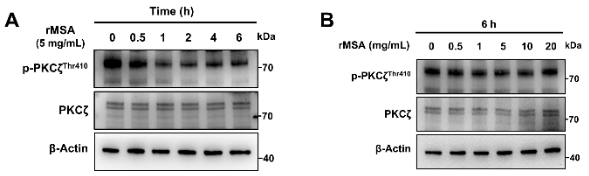


**Figure S13. The phosphorylation level of PKCζ remains unchanged after albumin treatment.**

Representative immunoblots of phospho-/total PKCζ in NMCMs treated with rMSA over a time course (A), and a concentration gradient for 6 h (B). β-Actin served as the loading control.

**Figure S14**

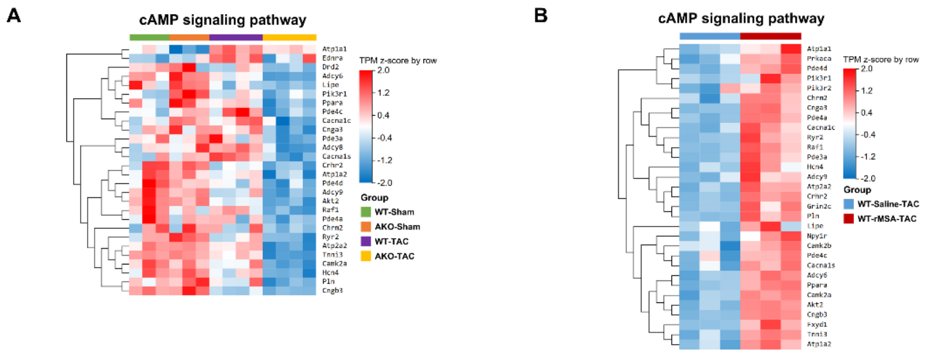


**Figure S14. Heatmaps of genes involved in cAMP signaling pathway.**

Heatmaps of cAMP signaling pathway transcriptomic profiles from cardiac tissues of WT-Sham, AKO-Sham, WT-TAC, and AKO-TAC groups, visualizing genes down-regulated in AKO-TAC (vs. WT-TAC) (A, n = 3–4); and from cardiac tissues of WT-Saline-TAC and WT-rMSA-TAC groups, visualizing genes up-regulated in WT-rMSA-TAC group (vs. WT-Saline-TAC) (B, n = 3).

#### Figure S15

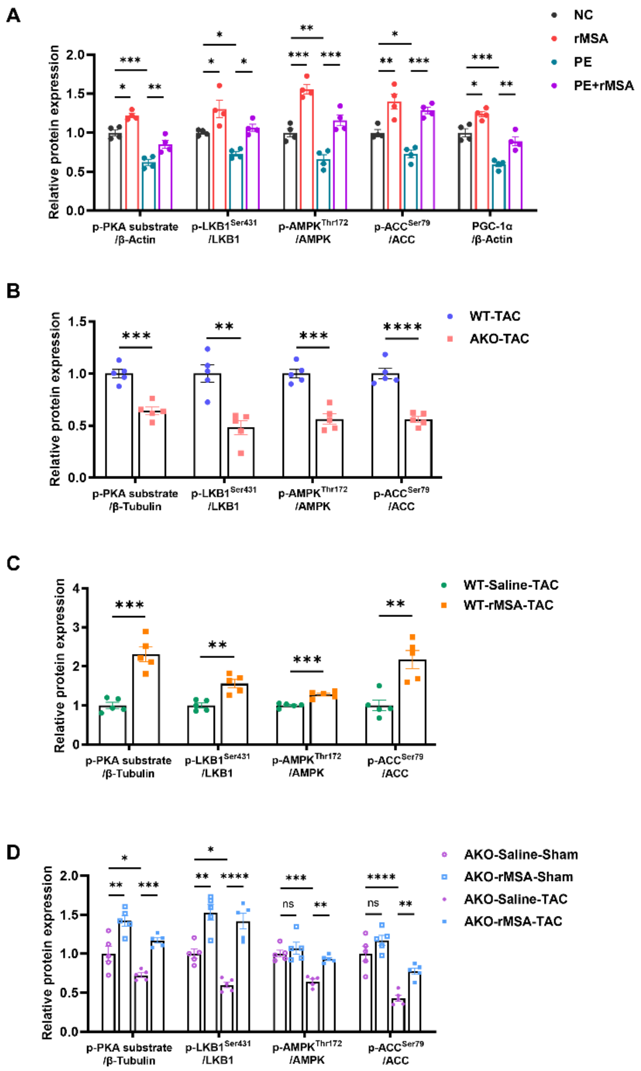


**Figure S15. Quantification of key PKA-LKB1-AMPK signaling components in the PE-induced cell model and TAC-induced mice with varied genotypes and treatment regimens.**

(A) Quantification of phosphorylated PKA substrate, phospho-/total LKB1, AMPK, ACC, and PGC-1α protein levels in the PE-induced cardiomyocyte hypertrophy model (n = 4).

(B–D) Quantification of phosphorylated PKA substrate, phospho-/total LKB1, AMPK, ACC, and PGC-1α protein levels in cardiac tissues from WT and AKO mice after TAC 6 weeks (B, n = 5), from WT mice treated with rMSA or saline after TAC 8 weeks (C, n = 5), from AKO mice treated with rMSA or saline after sham or TAC 8 weeks (D, n = 5).

Mean ± SEM, two-way ANOVA followed by Tukey's multiple comparisons test (A and D) or unpaired two-tailed Student's *t*-test (B and C). ns, not significant; *$p<0.05$, **$p<0.01$, ***$p<0.001$, ****$p<0.0001$.

**Figure S16**

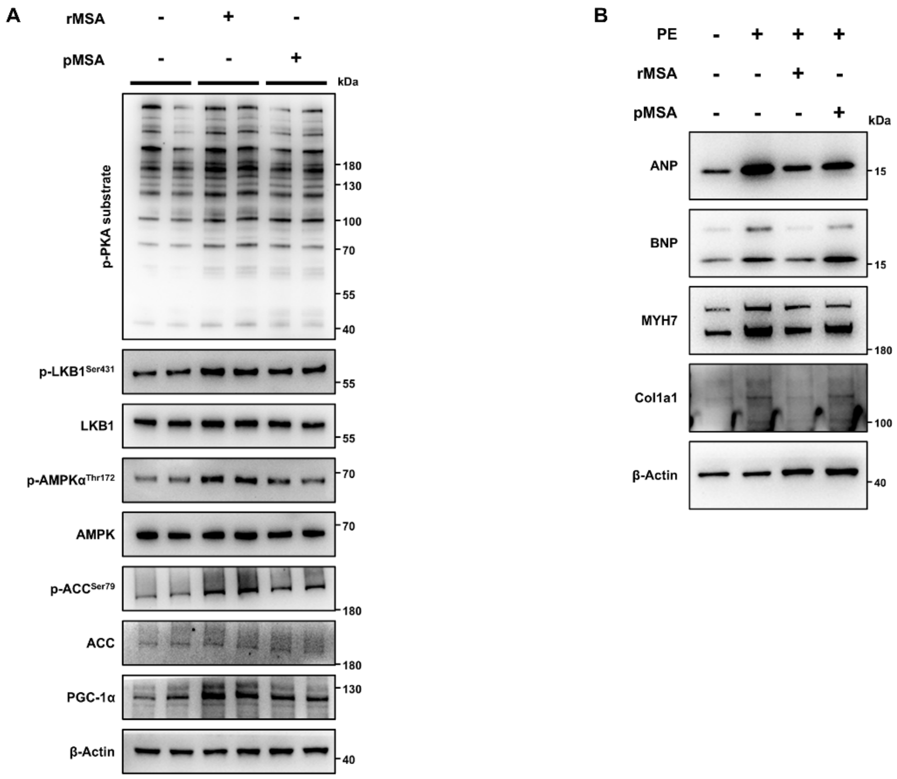


**Figure S16. rMSA more effectively activates the PKA-LKB1-AMPK signaling axis and attenuates PE-induced cardiomyocyte hypertrophy than plasma-derived mouse serum albumin (pMSA) in NMCMs.**

(A) Activation of the PKA-LKB1-AMPK signaling axis was assessed by representative immunoblots of phosphorylated PKA substrate, phospho-/total LKB1, AMPK, ACC, and PGC-1α protein levels in NMCMs following treatment with rMSA or pMSA (n = 4).

(B) Representative immunoblots of hypertrophy markers (ANP, BNP, MYH7, and Collagen I) in PE-induced NMCMs following treatment with rMSA or pMSA (n = 4).

(A and B) β-Actin served as the loading control.

**Table S1. Echocardiographic parameters in WT and AKO mice across ages.**

| | Young-8W | | Aged-24M | |
|---|---|---|---|---|
| | WT (n=6) | AKO (n=6) | WT (n=8) | AKO (n=8) |
| LVEF (%) | 66.23 ± 5.49 | 64.1 ± 3.94 | 49.35 ± 4.96$^{\#}$ | 44.81 ± 14.43 |
| LVFS (%) | 35.74 ± 4.13 | 34.23 ± 2.97 | 24.62 ± 2.86$^{\#}$ | 22.48 ± 8.38 |
| LV Vol;d (μL) | 47.74 ± 9.03 | 54.94 ± 10.69 | 66.54 ± 16.1 | 70.21 ± 23.53 |
| LV Vol;s (μL) | 16.21 ± 4.5 | 19.55 ± 3.41 | 34.12 ± 10.29 | 39.87 ± 19.18 |
| LVID;d (mm) | 3.4 ± 0.26 | 3.6 ± 0.29 | 3.89 ± 0.42 | 3.96 ± 0.54 |
| LVID;s (mm) | 2.19 ± 0.23 | 2.36 ± 0.16 | 2.94 ± 0.38$^{\#}$ | 3.08 ± 0.63 |
| LVPW;d (mm) | 0.78 ± 0.06 | 0.77 ± 0.07 | 1.1 ± 0.29 | 1.02 ± 0.31 |
| LVPW;s (mm) | 1.22 ± 0.11 | 1.17 ± 0.05 | 1.3 ± 0.3 | 1.2 ± 0.37 |
| LVAW;d (mm) | 0.76 ± 0.11 | 0.88 ± 0.08 | 1.26 ± 0.31$^{\#}$ | 1.22 ± 0.32 |
| LVAW;s (mm) | 1.18 ± 0.13 | 1.31 ± 0.13 | 1.65 ± 0.27$^{\#}$ | 1.55 ± 0.36 |
| LV Mass (mg) | 85.61 ± 9.51 | 103.46 ± 15.83 | 196.46 ± 52.88$^{\#}$ | 187.17 ± 48.9 |
| LV Mass Cor (mg) | 68.49 ± 7.61 | 82.77 ± 12.66 | 157.17 ± 42.3$^{\#}$ | 149.73 ± 39.12 |

All data are expressed as mean ± standard deviation. $^{\#}p<0.05$ vs. WT-Young-8W.

**Table S2. Echocardiographic parameters in WT and AKO mice 2 weeks post-TAC.**

| | Sham | | TAC-2W | |
|---|---|---|---|---|
| | WT (n=6) | AKO (n=6) | WT (n=9) | AKO (n=11) |
| LVEF (%) | 63.16 ± 6.37 | 62.3 ± 7.05 | 52.67 ± 6$^{\#}$ | 44.11 ± 6.59$^{\$}$ |
| LVFS (%) | 33.33 ± 4.76 | 33.2 ± 4.89 | 26.44 ± 3.64$^{\#}$ | 21.59 ± 3.92 |
| LV Vol;d (μL) | 39.91 ± 7.23 | 53.32 ± 12.57 | 50.41 ± 20.36 | 64.2 ± 14.73 |
| LV Vol;s (μL) | 14.69 ± 3.54 | 20.8 ± 8.92 | 24.43 ± 12.22 | 35.96 ± 9.5 |
| LVID;d (mm) | 3.15 ± 0.23 | 3.55 ± 0.34 | 3.43 ± 0.56 | 3.83 ± 0.39 |
| LVID;s (mm) | 2.1 ± 0.21 | 2.38 ± 0.4 | 2.53 ± 0.49 | 3.01 ± 0.35$^{\$}$ |
| LVPW;d (mm) | 0.88 ± 0.04 | 0.89 ± 0.19 | 1.15 ± 0.34 | 0.9 ± 0.18 |
| LVPW;s (mm) | 1.13 ± 0.13 | 1.19 ± 0.12 | 1.46 ± 0.34$^{\#}$ | 1.12 ± 0.13$^{\$}$ |
| LVAW;d (mm) | 1.32 ± 0.16 | 1.17 ± 0.18 | 1.26 ± 0.28 | 1.2 ± 0.27 |
| LVAW;s (mm) | 1.82 ± 0.13 | 1.68 ± 0.27 | 1.67 ± 0.32 | 1.59 ± 0.25 |
| LV Mass (mg) | 128.62 ± 13.55 | 141.05 ± 32.55 | 168.05 ± 38.38$^{\#}$ | 159.64 ± 15.23 |
| LV Mass Cor (mg) | 102.89 ± 10.84 | 112.84 ± 26.04 | 134.44 ± 30.71$^{\#}$ | 127.71 ± 12.18 |

All data are expressed as mean ± standard deviation. $^{\#}p<0.05$ vs. WT-Sham. $^{\$}p<0.05$ vs. WT-TAC-2W.

**Table S3. Echocardiographic parameters in WT and AKO mice 6 weeks post-TAC.**

| | Sham | | TAC-6W | |
|---|---|---|---|---|
| | WT (n=7) | AKO (n=7) | WT (n=9) | AKO (n=9) |
| LVEF (%) | 63.69 ± 8.25 | 63.95 ± 5.01 | 43.98 ± 8.92$^{\#}$ | 31.81 ± 7.03$^{\$}$ |
| LVFS (%) | 34.01 ± 5.88 | 33.81 ± 3.75 | 21.63 ± 5.04$^{\#}$ | 15.01 ± 3.68$^{\$}$ |
| LV Vol;d (μL) | 47.87 ± 14.06 | 34.26 ± 3.79 | 73.49 ± 19.68$^{\#}$ | 93.19 ± 25.55 |
| LV Vol;s (μL) | 17.65 ± 7.48 | 12.31 ± 1.8 | 42.36 ± 16.78$^{\#}$ | 64.42 ± 22.35$^{\$}$ |
| LVID;d (mm) | 3.38 ± 0.41 | 2.97 ± 0.13 | 4.05 ± 0.47$^{\#}$ | 4.48 ± 0.53 |
| LVID;s (mm) | 2.24 ± 0.37 | 1.96 ± 0.12 | 3.19 ± 0.53$^{\#}$ | 3.82 ± 0.56$^{\$}$ |
| LVPW;d (mm) | 0.85 ± 0.14 | 0.99 ± 0.19 | 1.02 ± 0.22 | 0.99 ± 0.15 |
| LVPW;s (mm) | 1.31 ± 0.19 | 1.43 ± 0.23 | 1.25 ± 0.21 | 1.13 ± 0.15 |
| LVAW;d (mm) | 0.91 ± 0.18 | 0.96 ± 0.06 | 1.13 ± 0.13$^{\#}$ | 1.06 ± 0.15 |
| LVAW;s (mm) | 1.3 ± 0.22 | 1.36 ± 0.08 | 1.49 ± 0.1 | 1.37 ± 0.16 |
| LV Mass (mg) | 102.42 ± 15.6 | 98.63 ± 18.73 | 182.47 ± 39.18$^{\#}$ | 198.55 ± 34.17 |
| LV Mass Cor (mg) | 81.93 ± 12.48 | 78.9 ± 14.99 | 145.98 ± 31.35$^{\#}$ | 158.84 ± 27.34 |

All data are expressed as mean ± standard deviation. $^{\#}p<0.05$ vs. WT-Sham. $^{\$}p<0.05$ vs. WT-TAC-6W.

**Table S4. Echocardiographic parameters in WT mice administrated with rMSA and saline 8 weeks post-TAC.**

| | Sham | | TAC-8W | |
|---|---|---|---|---|
| | WT-Saline (n=7) | WT-rMSA (n=7) | WT-Saline (n=14) | WT-rMSA (n=12) |
| LVEF (%) | 68.58 ± 4.49 | 70.86 ± 7.96 | 47.86 ± 9.09$^{\#}$ | 61.99 ± 9.51$^{\$}$ |
| LVFS (%) | 37.69 ± 3.83 | 39.71 ± 6.51 | 23.97 ± 5.5$^{\#}$ | 33.32 ± 6.86$^{\$}$ |
| LV Vol;d (μL) | 53.79 ± 13.38 | 45.64 ± 10.69 | 71.13 ± 11.76$^{\#}$ | 64.49 ± 12.92 |
| LV Vol;s (μL) | 16.94 ± 5.21 | 13.32 ± 5.59 | 37.73 ± 10.91$^{\#}$ | 25.31 ± 10.41$^{\$}$ |
| LVID;d (mm) | 3.56 ± 0.41 | 3.32 ± 0.35 | 4.01 ± 0.28$^{\#}$ | 3.85 ± 0.33 |
| LVID;s (mm) | 2.21 ± 0.29 | 2 ± 0.31 | 3.06 ± 0.39$^{\#}$ | 2.58 ± 0.44$^{\$}$ |
| LVPW;d (mm) | 0.78 ± 0.15 | 0.72 ± 0.17 | 1.01 ± 0.14 | 0.97 ± 0.27 |
| LVPW;s (mm) | 1.22 ± 0.26 | 1.23 ± 0.28 | 1.26 ± 0.17 | 1.27 ± 0.39 |
| LVAW;d (mm) | 0.92 ± 0.23 | 0.85 ± 0.21 | 1.03 ± 0.17 | 0.89 ± 0.17 |
| LVAW;s (mm) | 1.36 ± 0.18 | 1.35 ± 0.17 | 1.42 ± 0.18 | 1.39 ± 0.15 |
| LV Mass (mg) | 104.64 ± 17.79 | 84.3 ± 8.27 | 166.47 ± 32.72$^{\#}$ | 134.31 ± 24.16$^{\$}$ |
| LV Mass Cor (mg) | 83.72 ± 14.23 | 67.44 ± 6.62 | 133.18 ± 26.17$^{\#}$ | 107.45 ± 19.33$^{\$}$ |

All data are expressed as mean ± standard deviation. $^{\#}p<0.05$ vs. WT-Saline-Sham. $^{\$}p<0.05$ vs. WT-Saline-TAC-8W.

**Table S5. Echocardiographic parameters in AKO mice administrated with rMSA and saline 8 weeks post-TAC.**

| | Sham | | TAC-8W | |
|---|---|---|---|---|
| | AKO-Saline (n=7) | AKO-rMSA (n=7) | AKO-Saline (n=8) | AKO-rMSA (n=8) |
| LVEF (%) | 59.64 ± 5.17 | 60.74 ± 5.53 | 30.4 ± 1.68$^{\#}$ | 38.73 ± 4.61$^{\$}$ |
| LVFS (%) | 31.2 ± 3.56 | 31.83 ± 3.74 | 14.14 ± 0.89$^{\#}$ | 18.48 ± 2.51$^{\$}$ |
| LV Vol;d (μL) | 59.92 ± 9.27 | 52.87 ± 10.66 | 78.57 ± 15.39$^{\#}$ | 65.21 ± 6.85 |
| LV Vol;s (μL) | 24.41 ± 5.97 | 21.23 ± 6.56 | 54.62 ± 10.51$^{\#}$ | 39.92 ± 4.69$^{\$}$ |
| LVID;d (mm) | 3.74 ± 0.24 | 3.54 ± 0.31 | 4.18 ± 0.36$^{\#}$ | 3.88 ± 0.17 |
| LVID;s (mm) | 2.58 ± 0.26 | 2.42 ± 0.33 | 3.59 ± 0.3$^{\#}$ | 3.16 ± 0.16$^{\$}$ |
| LVPW;d (mm) | 0.86 ± 0.15 | 0.9 ± 0.13 | 0.92 ± 0.28 | 0.85 ± 0.13 |
| LVPW;s (mm) | 1.23 ± 0.2 | 1.3 ± 0.26 | 1.02 ± 0.2 | 1.07 ± 0.21 |
| LVAW;d (mm) | 0.91 ± 0.15 | 1.17 ± 0.24 | 1 ± 0.12 | 1.11 ± 0.24 |
| LVAW;s (mm) | 1.34 ± 0.17 | 1.57 ± 0.26 | 1.18 ± 0.12 | 1.4 ± 0.19 |
| LV Mass (mg) | 120.73 ± 15.97 | 139.41 ± 26.5 | 160.97 ± 26.28$^{\#}$ | 149.36 ± 27.58 |
| LV Mass Cor (mg) | 96.58 ± 12.77 | 111.53 ± 21.2 | 128.78 ± 21.02$^{\#}$ | 119.49 ± 22.06 |

All data are expressed as mean ± standard deviation. $^{\#}p<0.05$ vs. AKO-Saline-Sham. $^{\$}p<0.05$ vs. AKO-Saline-TAC-8W.

**Table S6. Primer sequences used in this study.**

| Gene | Forward (5'-3') | Reverse (5'-3') |
|---|---|---|
| *Mouse-Nppa* | GTGCGGTGTCCAACACAGAT | TCCAATCCTGTCAATCCTACCC |
| *Mouse-Nppb* | GAGGTCACTCCTATCCTCTGG | GCCATTTCCTCCGACTTTTCTC |
| *Mouse-Myh7* | ACTGTCAACACTAAGAGGGTCA | TTGGATGATTTGATCTTCCAGGG |
| *Mouse-Col1a1* | GCTCCTCTTAGGGGCCACT | CCACGTCTCACCATTGGGG |
| *Mouse-Col3a1* | ACGTAGATGAATTGGGATGCAG | GGGTTGGGGCAGTCTAGTG |
| *Mouse-Ctgf* | GGGCCTCTTCTGCGATTTC | ATCCAGGCAAGTGCATTGGTA |
| *Mouse-Tgfb1* | CTCCCGTGGCTTCTAGTGC | GCCTTAGTTTGGACAGGATCTG |
| *Mouse-Tnfa* | CCCCAAAGGGATGAGAAGTT | ACTTGGTGGTTTGCTACGA |
| *Mouse-Il6* | GATAAGCTGGAGTCACAGAAGG | TTGCCGAGTAGATCTCAAAGT |
| *Mouse-Mcp1* | CCACTCACCTGCTGCTACTCAT | TGGTGATCCTCTTGTAGCTCTCC |
| *Mouse-Ppara* | AGAGCCCCATCTGTCCTCTC | ACTGGTAGTCTGCAAAACCAAA |
| *Mouse-Ppargc1a* | TATGGAGTGACATAGAGTGTGCT | CCACTTCAATCCACCCAGAAAG |
| *Mouse-Cpt1b* | GCACACCAGGCAGTAGCTTT | CAGGAGTTGATTCCAGACAGGTA |
| *Mouse-Slc25a20* | GACGAGCCGAAACCCATCAG | AGTCGGACCTTGACCGTGT |
| *Mouse-Acadm* | AGGGTTTAGTTTTGAGTTGACGG | CCCCGCTTTTGTCATATTCCG |
| *Mouse-Acadl* | TCTTTTCCTCGGAGCATGACA | GACCTCTCTACTCACTTCTCCAG |
| *Mouse-Acadvl* | CTACTGTGCTTCAGGGACAAC | CAAAGGACTTCGATTCTGCCC |
| *Mouse-Acaa2* | CTGCTACGAGGTGTGTTCATC | AGCTCTGCATGACATTGCCC |
| *Mouse-Cd36* | ATGGGCTGTGATCGGAACTG | TTTGCCACGTCATCTGGGTTT |
| *Mouse-Slc27a1* | CTGGGACTTCCGTGGACCT | TCTTGCAGACGATACGCAGAA |
| *Mouse-Acsl1* | TGCCAGAGCTGATTGACATTC | GGCATACCAGAAGGTGGTGAG |
| *Mouse-Errα* | CCAGGCTTCTCCTCACTGTC | GCCCCCTCTTCATCTAGGAC |

| Gene | Forward (5'-3') | Reverse (5'-3') |
|---|---|---|
| *Mouse-Nrf1* | AGAAACGGAAACGGCCTCAT | ACTCATCCAACGTGGCTCTG |
| *Mouse-Ppargc1b* | TCTGACGTGGACGAGCTTTCA | CCCGTCCTTCAGAGCGTCAG |
| *Mouse-Tfam* | CCCGGCAGAGACGGTTAAAA | TCCCTGAGCCGAATCATCCT |
| *Mouse-Drp1* | ACAACAGGAGAAGAAAATGGAGT | GATGGATTGGCTCAGGGCTT |
| *Mouse-Fis1* | TGTCCAAGAGCACGCAATTTG | CCTCGCACATACTTTAGAGCCTT |
| *Mouse-Mff* | ATGCCAGTGTGATAATGCAAGT | CTCGGCTCTCTTCGCTTTG |
| *Mouse-Opa1* | TGGAAAATGGTTCGAGAGTCAG | CATTCCGTCTCTAGGTTAAAGCG |
| *Mouse-Mfn1* | CCTACTGCTCCTTCTAACCCA | AGGGACGCCAATCCTGTGA |
| *Mouse-Mfn2* | AGAACTGGACCCGGTTACCA | CACTTCGCTGATACCCCTGA |
| *Mouse-Ndufv1* | TGGATCGCTCGACAGACATT | GCACTTGTTTATTGTCCAGTTTCCC |
| *Mouse-Ndufs1* | CCAGGAGGTTCTTGCTGACC | TGTCTCAGCATATGGACGGC |
| *Mouse-Sdhc* | TCCAGGCCGGAACTCAAGAT | CAGCCAGACCTGGGGTATTG |
| *Mouse-Sdha* | GGAACACTCCAAAAACAGACCT | CCACCACTGGGTATTGAGTAGAA |
| *Mouse-Uqcrfs1* | GAGCCACCTGTTCTGGATGTG | GCACGACGATAGTCAGAGAAGTC |
| *Mouse-Uqcrc1* | GAGGTGACTCGGGGCAAAAA | GGAAGGACAACACAGTTCTCAAT |
| *Mouse-Cox7b* | TGGAACATGTCCCCTGTTGG | CACCATGCTTGGGCTTCATT |
| *Mouse-Cox4i1* | CAGCCTTTCCAGGGATGAGAA | AGAAGCCCTGAATGGGGTTG |
| *Mouse-Atp5o* | ATCTGTGGTCAGGCCCTTTG | CCAGAGACACTTTGGGGTCC |
| *Mouse-Atp5a1* | TCTCCATGCCTCTAACACTCG | CCAGGTCAACAGACGTGTCAG |
| *Mouse-Pink1* | CCCACACCCTAACATCATCC | ACTGGGAGTCTGCTCCTCAA |
| *Mouse-Prkn* | GGGATTCAGAAGCAGCCAGA | CAGCAAGATGGGCCCTGG |
| *Mouse-Actb* | GGCTGTATTCCCCTCCATCG | CCAGTTGGTAACAATGCCATGT |
| *Mouse-Alb* | TGCTTTTCCAGGGGTGTGTT | TTACTTCCTGCACTAATTTGGCA |

| Gene | Forward (5'-3') | Reverse (5'-3') |
|---|---|---|
| *Mouse-Hk2* | GCCAGCCTCTCCTGATTTTAGTGT | GGGAACACAAAAGACCTCTTCTGG |
| *Mouse-mtNd1* | ATCTTATCCACGCTTCCGTTACG | TATGGTGGTACTCCCGCTGTA |

**Table S7. Primary and secondary antibodies used in this study.**

| Antibody | Product code | Source | Dilution ratio |
|---|---|---|---|
| Albumin | ab207327 | Abcam | 1:2000 |
| Transferrin | 17435-1-AP | Proteintech | 1:1000 |
| PGC-1α | 66369-1-Ig | Proteintech | 1:5000 |
| TFAM | 22586-1-AP | Proteintech | 1:5000 |
| β-Tubulin | AC021 | ABclonal | 1:5000 |
| PPARα | 66826-1-Ig | Proteintech | 1:2000 |
| NDUFS1 | 12444-1-AP | Proteintech | 1:2000 |
| SDHA | 66588-1-Ig | Proteintech | 1:5000 |
| UQCRC1 | 67888-1-Ig | Proteintech | 1:5000 |
| COX IV | ab14744 | Abcam | 1:5000 |
| ATP5O | 10994-1-AP | Proteintech | 1:1000 |
| ATP8 | 26723-1-AP | Proteintech | 1:1000 |
| PINK1 | 23274-1-AP | Proteintech | 1:1000 |
| Parkin | 14060-1-AP | Proteintech | 1:1000 |
| LC3B | 83506 | CST | 1:1000 |
| ANP | 27426-1-AP | Proteintech | 1:2000 |
| BNP | A2179 | ABclonal | 1:1000 |
| Collagen Type I | 67288-1-Ig | Proteintech | 1:5000 |
| β-Actin | TA-09 | ZSGB-Bio | 1:2000 |
| p-PKA Substrate | 9624S | CST | 1:1000 |
| p-LKB1 (Ser428) | 3482S | CST | 1:1000 |
| LKB1 | 3047S | CST | 1:1000 |
| p-AMPKα (Thr172) | 2535S | CST | 1:1000 |
| AMPKα | 2532S | CST | 1:1000 |
| p-ACC (Ser79) | 11818S | CST | 1:1000 |
| ACC | 3676S | CST | 1:1000 |
| CAMKK2 | 11549-1-AP | Proteintech | 1:1000 |
| p-PKCζ/λ (Thr410/403) | 9378T | CST | 1:1000 |

| Antibody | Product code | Source | Dilution ratio |
|---|---|---|---|
| PKCζ | 9368T | CST | 1:1000 |
| α-Actinin | A7811 | Sigma | 1:500 |
| Albumin (for IF) | 16475-1-AP | Proteintech | 1:200 |
| HRP-linked mouse IgG | 7076S | CST | 1:3000 |
| HRP-linked rabbit IgG | 7074S | CST | 1:3000 |
| TRITC-conjugated goat anti-rabbit IgG | ZF-0316 | ZSGB-Bio | 1:200 |
| FITC-conjugated goat anti-rabbit IgG | ZF-0311 | ZSGB-Bio | 1:200 |
| TRITC-conjugated goat anti-mouse IgG | ZF-0313 | ZSGB-Bio | 1:200 |
| FITC-conjugated goat anti-mouse IgG | ZF-0312 | ZSGB-Bio | 1:200 |
